# A Multiplexed Bacterial Two-Hybrid for Rapid Characterization of Protein-Protein Interactions and Iterative Protein Design

**DOI:** 10.1101/2020.11.12.377184

**Authors:** W. Clifford Boldridge, Ajasja Ljubetič, Hwangbeom Kim, Nathan Lubock, Dániel Szilágyi, Jonathan Lee, Andrej Brodnik, Roman Jerala, Sriram Kosuri

## Abstract

Myriad biological functions require protein-protein interactions (PPIs), and engineered PPIs are crucial for applications ranging from drug design to synthetic cell circuits. Understanding and engineering specificity in PPIs is particularly challenging as subtle sequence changes can drastically alter specificity. Coiled-coils are small protein domains that have long served as a simple model for studying the sequence-determinants of specificity and have been used as modular building blocks to build large protein nanostructures and synthetic circuits. Despite their simple rules and long-time use, building large sets of well-behaved orthogonal pairs that can be used together is still challenging because predictions are often inaccurate, and, as the library size increases, it becomes difficult to test predictions at scale. To address these problems, we first developed a method called the Next-Generation Bacterial Two-Hybrid (NGB2H), which combines gene synthesis, a bacterial two-hybrid assay, and a high-throughput next-generation sequencing readout, allowing rapid exploration of interactions of programmed protein libraries in a quantitative and scalable way. After validating the NGB2H system on previously characterized libraries, we designed, built, and tested large sets of orthogonal synthetic coiled-coils. In an iterative set of experiments, we assayed more than 8,000 PPIs, used the dataset to train a novel linear model-based coiled-coil scoring algorithm, and then characterized nearly 18,000 interactions to identify the largest set of orthogonal PPIs to date with twenty-two on-target interactions.

## Introduction

Protein-protein interactions (PPIs) are integral to most biological functions and are required for such diverse processes as cell division, signaling, metabolism and transcription and translation^1^. Our ability to design and build functions and structures as complex as nature is still in its infancy, but is developing with advances in both protein design algorithms and gene synthesis capacities. For example, orthogonal *de novo* designed proteins and the careful reuse of well-characterized orthogonal interactions found in natural systems facilitate building nanoscale superstructures for applications in biology, biological engineering and materials science^2^. Supramolecular protein designs can be created using simple, natural protein families like coiled-coils, which have been used to build numerous designed protein assemblies^3–5^. However, identifying orthogonal natural proteins is difficult, as evolutionarily related proteins often display significant cross-interactions. Another method is to computational design *de novo* proteins; in particular, Rosetta-based designs have produced homodimers^6,7^ and heterodimers^8^. However, predicting orthogonal binding and designing large orthogonal sets remains beyond current de-novo design methods^3^.

Coiled-coils in particular have many useful characteristics for atomically precise designs of macromolecular structures. They are small, precisely oriented, and numerous sequence-based and computational models exist to describe their properties. First identified at the dawn of molecular biology by both Pauling^9^ and Crick^10^, coiled-coils are defined by their heptad repeat H-P-P-H-P-P-P (H = hydrophobic residue, P = polar residue). This relatively simple structure has given rise to many computational models to describe coiled-coil interactions, from the parametric Crick equations in 1953^11^ to contemporary linear models^12–15^. However, because of their shared similar structure, building large sets of orthogonally interacting coiled-coils, where all on-target interactions occur to the exclusion of all off-target interactions, is still difficult. Though numerous groups have attempted to create orthogonal sets of coiled-coils, they’ve been limited in size and still display significant off-target interactions^15,16^. Increasing our ability to design, build and characterize large sets of interacting proteins could help solve this problem by providing empirical data to improve computational models of PPIs. Simultaneously this would vastly increase the number of available orthogonal building blocks for nanoscale structural design allowing the creation of previously unbuildable structures.

Here we combine gene synthesis, a novel assay that allows for multiplexed bimolecular interaction screening, and a computational pipeline to design large libraries of orthogonally interacting coiled-coils. We first built and validated a novel assay, the Next-Generation Bacterial Two-Hybrid (NGB2H) system that has a number of unique advantages over other methodologies for characterizing protein libraries. In particular, the NGB2H system allows for screening of bimolecular interactions without having to test all-against-all libraries, direct large-scale synthesis using oligonucleotide arrays to explore design space, quantitative readouts on the entire library including negative interactions, and allows for understanding low affinity interactions inside the crowded cellular context. We did this iteratively with synthetically-designed libraries increasing in size from 256 interactions to more than 18,000 interactions. From this we identified the largest sets of orthogonal proteins to date and developed an improved coiled-coil design algorithm for future design purposes of this versatile protein domain.

## Results

### NGB2H system design

To allow iterative protein design using synthetic constructs, we built a generalizable, scalable bacterial two-hybrid system using a significantly modified version of the *B. pertussis* adenylate cyclase two-hybrid^17^ (Figure 1A, Supplementary Information Section 1). Briefly, the two-hybrid functions much as in Karimova et al.^17^, where interacting hybrid proteins reconstitute adenylate cyclase to produce cAMP which drives reporter gene expression. We measured relative transcription of a uniquely identifying DNA barcode residing in the reporter gene, which serves as a measure for interaction strength. The barcode is mapped to the two fully sequenced hybrid proteins at an early cloning step using high-throughput sequencing when the barcode and proteins are physically adjacent. This unambiguously identifies even highly homologous proteins and separates synthetic errors from programmed designs. Thus, measuring the relative barcode transcription provides a quantitative, massively multiplexed characterization of PPIs with short read sequencing. Because the NGB2H system uses a mapping step, it can use gene synthesis, rather than preconstructed libraries to create diversity, which further frees it from one-against-all or all-against-all testing common in two-hybrids. We made a number of other improvements, including: (1) titratable and inducible control of hybrid protein expression and optimized reporter response on a single plasmid, (2) a background strain with linear cAMP accumulation, (3) a GFP reporter instead of beta-galactosidase for more rapid individual characterization, (4) the use of multiple barcodes per construct to achieve better statistical certainty, and (5) a scarless cloning scheme that allows for library creation with any designed sequence. (More information Supplementary Information Section 1).

**Figure 1).**
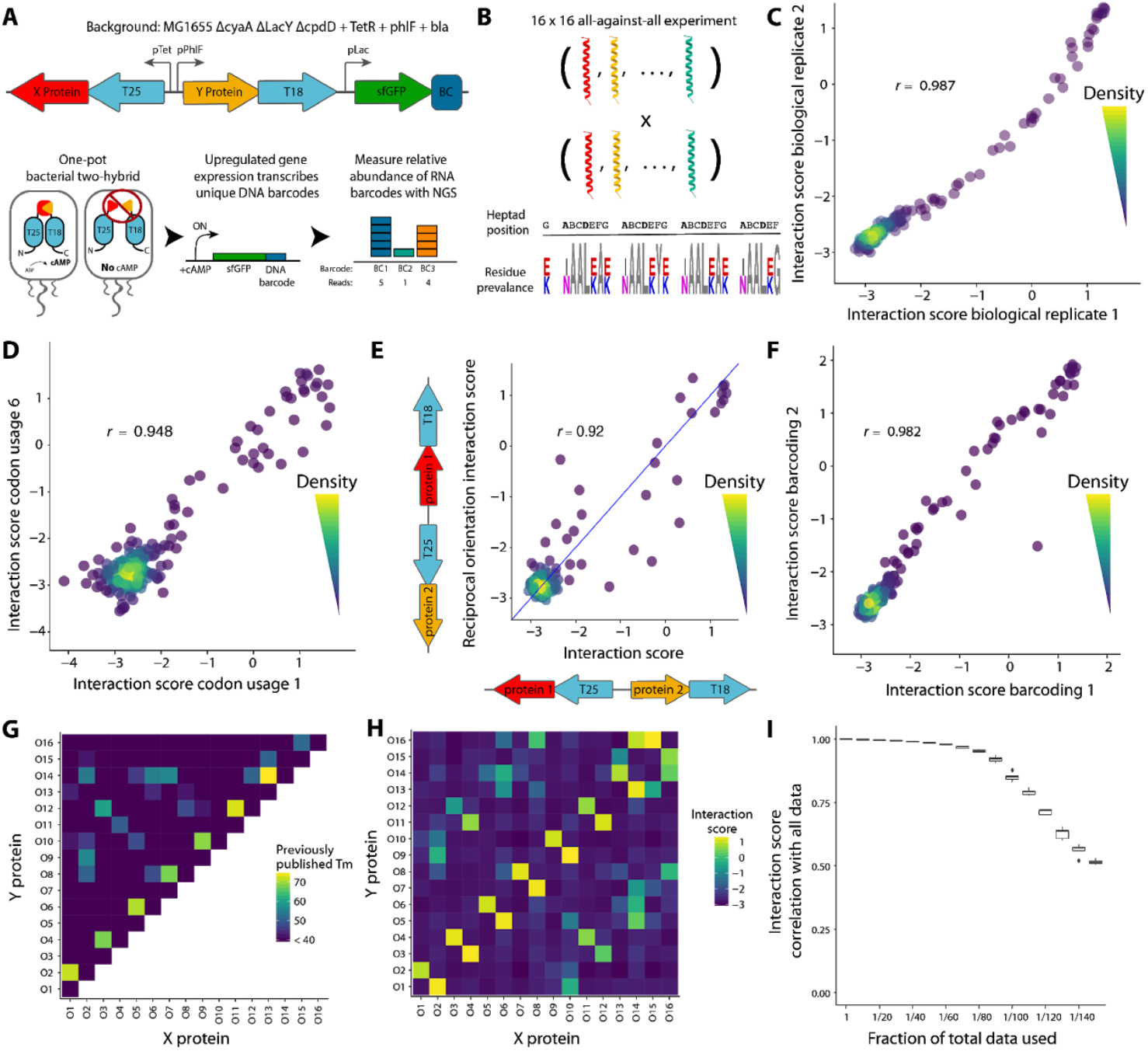
Design and validation of the NGB2H assay. A) (Top) Schematic of the NGB2H system reporter construct. T25, T18 - adenylate cyclase halves; BC - unique DNA barcode identifying the protein pair. (Bottom) Workflow of NGB2H system. Interacting proteins reconstitute adenylate cyclase, producing cAMP which drives gene expression of the barcoded sfGFP reporter. Relative barcode abundance is quantified using next generation sequencing (NGS). B) The CC0 Library is composed of 16 coiled-coils tested against one another. (Bottom) Sequence logo representing the diversity represented in the CC0 Library. Residues that vary are shown in color. C) Interaction scores of CC0 Library members are consistent between biological replicates (Pearson’s r > 0.98). D) Two different codon usages have consistent interaction scores (Pearson’s r > 0.94, representative sample). E) Interaction strength is similar (Pearson’s r > 0.92) regardless of which protein is attached to which half of adenylate cyclase. The blue line represents y = x. F) Interaction scores of separately barcoded, cloned and tested replicates are consistent (Pearson’s r > 0.98). G) Published circular dichroism (CD) melting point (Tm) data. H) Experimentally determined interaction scores. I) CC0 Library raw data can be subsampled and still correlate well with the full dataset. Boxplots represent the interquartile range of 50 random subsamples of the full data.

### Validation of the NGB2H system

After optimizing the system with single construct GFP measurements (Figure S1), we validated the NGB2H system with 256 previously characterized interactions^15^, which we call the CC0 Library. The CC0 Library is a set of sixteen *de novo* designed, orthogonal, heterodimeric coiled-coils tested in an all-against-all configuration. The proteins are highly similar, four heptad coiled-coils which vary only at the a-position (Ile/Asn), e-position and g-position (Lys/Glu) (Figure 1B). We designed the CC0 Library to be compatible with our system (Figure S2A), barcoded and cloned it (Figure S3A, S4). After inducing the two-hybrid for six hours, we took samples for RNA and DNA extraction to measure interaction strength and normalize for plasmid abundance, respectively. We obtained high-quality measurements for all 256 protein pairs and calculated an Interaction score where

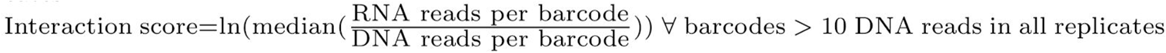

The NGB2H assay was highly replicable, with biological replicates having similar Interaction scores (Pearson’s *r* > 0.98, p < 10^−15^) with a dynamic range of more than 100-fold (Figure 1C).

We checked several internal controls to validate the measurements of the NGB2H assay. First, as the protein code is degenerate, we screened nine different codon usages for each pair of proteins. Different codon usages showed consistent Interaction scores (representative pair Figure 1D) with all usages correlating with Pearson’s *r* > 0.92 and p < 10^−15^ (Figure S5), demonstrating minimal effects from DNA sequence variation and low levels of noise in Interaction scores.

We also compared the Interaction scores of protein pairs when attached to the other half of the two-hybrid, which we call the reciprocal orientation. We found that the CC0 Library has a strong correlation between the primary and reciprocal orientations (Pearson’s *r* = 0.92, p < 10^−15^) (Figure 1E), indicating that the biological machinery of the NGB2H system faithfully recapitulates the biochemical interaction. In addition, a portion of our library contained frameshift mutations which should not create functional PPIs. As expected, Interaction scores of constructs with indels cluster at the bottom of the range of correct constructs (Figure S6). Lastly, to show that the NGB2H system does not suffer from barcode effects or selection pressure from the repeated cloning steps, we replicated the assay with an independent re-barcoding and re-cloning of the CC0 Library which showed strong correlation with the first iteration’s Interaction scores (Pearson’s *r* > 0.98, p < 10^−15^) (Figure 1F).

Having confirmed the internal consistency of the CC0 Library, we compared it to the previously published results. When compared to circular dichroism data published in Crooks et al.^15^, we found the NGB2H system’s dynamic range correlated well with melting temperatures greater than 40°C (Figure 1G, 1H). Given the differences in technique – *in vivo* versus *in vitro*, interaction strength versus helicity – the correlation between the Interaction score and Tm (Pearson’s *r* > 0.75, p < 10^−15^, Figure S7) largely validates the NGB2H system. Finally, the NGB2H system needs to be highly scalable. To test its scalability, we downsampled the raw sequencing reads between 10 and 150-fold and found strong agreement with our full dataset even when downsampled 100-fold (Pearson’s *r* > 0.85, p < 10^−15^, Figure 1I), which implies the ability to accurately screen ~25,000 interactions with an equal number of reads.

### Computational design of large sets of orthogonal coiled-coils

To computationally predict large, orthogonal sets of coiled-coils for empirical verification, we built a two-step computational pipeline (Figure 2A). In brief, we calculated 16.7 million interaction scores for all four heptad coiled-coils with Ile or Asn at the a-position and Glu or Lys at the e- and g-positions using the scoring model from Potapov et al^13^. We then identified sets with an orthogonality gap—those sets where all on-target interactions had a higher predicted score than all off-target interactions. Though computationally challenging, this is tractable as a variant of the maximum independent set problem^18^. We identified the fifteen largest sets and designed each of them with three different backbones (each containing different residues at the noncontact b, c, and f-positions) to investigate their contribution to coiled-coil stability^19^. We combined these with two sets of controls spanning eleven backbones, for a total of 56 sets containing between 64 to 961 interactions (8,169 interactions overall), which we named the CCNG1 Library. After testing a subset of the CCNG1 Library to validate our in-house designs, (Figure S8, S9, Supplementary Information Section 8.3), we designed (Figure S2C), cloned (Figure S3C, S4) and ran the NGB2H assay, from which we collected quality data (Figure S10) on 8,073 interactions.

**Figure 2).**
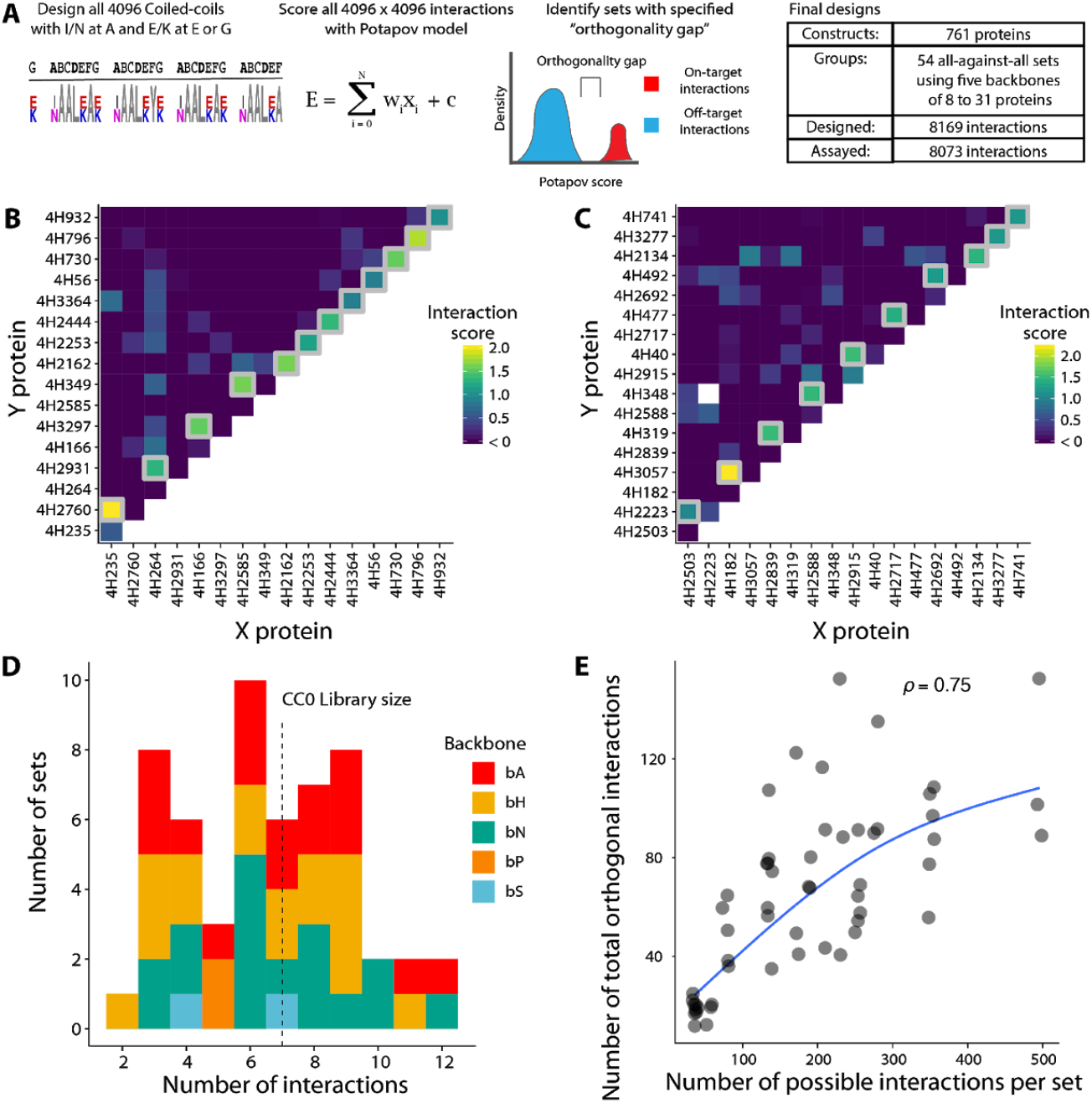
Large orthogonal sets of coiled-coils from the CCNG1 library. A) Schematic of the CCNG1 Library design. All 4-heptad coiled-coils with variation at positions a, e, and g were scored for interactions with the model from Popatov et al., and sets of coiled-coils with large orthogonality gaps were identified. In total we designed and tested 54 sets of orthogonal coiled-coils. B) The set of coiled-coils with the largest number of on-target orthogonal interactions (12 on-target interactions). Grey boxes identify on-target interactions. C) Set of orthogonal coiled-coils with the largest number of total (on-target and non-interacting) interactions (153 total interactions). Grey boxes represent on-target interactions. D) Number of interactions per set of coiled-coils. Dashed line represents the number of on-target orthogonal interactions in the CC0 library. Colors show different backbones used, while the interfacial residues stayed the same. E) Set size increases more rapidly than the number of pairs in the largest orthogonal subset. Blue line represents a spline with degree two.

### Large orthogonal sets in the CCNG1 library

Although we designed our coiled-coils to be orthogonal, the current state-of-the-art design algorithms are relatively inaccurate. Thus, similar to our designs, we reduced the problem to the maximum independent set problem to identify the largest orthogonal subset of each set. We were able to identify a set of orthogonal coiled-coils that contains twelve pairs, which includes four heterodimers and eight homodimers (Figure 2B). We have also identified a set of seven heterodimers and three homodimers (Figure 2C), that has fewer on-target interactions (ten versus twelve), but contains more proteins (seventeen versus sixteen) and therefore more total potential interactions (153 versus 136).

We characterized the number of on-target interactions across the CCNG1 Library and found 20 of our 51 sets have more than the seven on-target orthogonal interactions in the state-of-the-art set from Crooks et al., (Figure 2D), though many share the same interfacial residues (Figure S11). We found that these orthogonal sets were composed of between four and seventeen proteins (Figure S12), five of which are larger than the CC0 Library. As the CCNG1 Library represents the first large scale systematic investigation into the effects of variation at the b, c, and f-positions, we sought to understand how these positions influenced interactions. We tested six backbones containing the same interfacial residues as the CC0 Library (Figure S13, Supplementary Information Section 8.4) and found that charged backbones led to less specific interaction profiles. To understand the limits of producing orthogonal interactions within highly constrained sequence space, we compared the number of total pairs in each set (the sum of interacting and non-interacting pairs) to the number of pairs in its largest orthogonal subset (Figure 2E). We found that the number of orthogonal pairs appears to increase progressively slower as set size increased, suggesting much larger sets would need to be screened to identify proportionally more orthogonal coiled-coils or, alternatively, the accuracy of prediction algorithms would need to be improved.

### Improving the state-of-the-art coil-coiled interaction prediction algorithms

The CCNG1 Library dataset represents the largest dataset of coiled-coil interactions to date. We reasoned that our data could serve as a training set to improve on currently available models. To benchmark current models, we computed scores using algorithms bCipa^14^, Potapov/SVR^13^, Fong/SVM^20^ and Vinson/CE^12^ which are all linear models with features for amino acid pairings. Each algorithm is only weakly predictive of our measured interactions with the bA backbone (Figure 3A), as all models have an R^2^ < 0.2. Notably, each algorithm predicted the strongest interactions well, but also predicted many weak interactions that when measured had high Interaction scores. We built several linear models similar to bCipa which included numerous innovations (Supplementary Information Section 3). First, we trained a model on our data that only included weights for the a-, e- and g-position combinations. We also created versions of this simple model with terms for either consecutive residues in the a-position of the same protein or separate terms for weights at the N-terminal a-position, where fraying may occur (Figure S14A). We then expanded these models with a novel scoring technique, which we call heptad shifts (Figure S14B). In short, we expect the predominant form of coiled-coil interaction to be the alignment of heptads that has the strongest interaction, which is not necessarily all four heptads aligned from the N-terminus, but could be an interface of three or fewer heptads. All of our heptad shifting scoring algorithms were significantly better than the corresponding non-shifting versions and our N-terminal a-positions weights algorithm was significantly better than both the basic algorithm and the consecutive a-position algorithm (Figure 3B). Thus, our final model which we call iCipa uses heptad shifting and terms for the N-terminal a-positions, and it is more predictive of CCNG1 Interaction scores than previous models with an R^2^ = 0.27 (Figure 3C).

**Figure 3).**
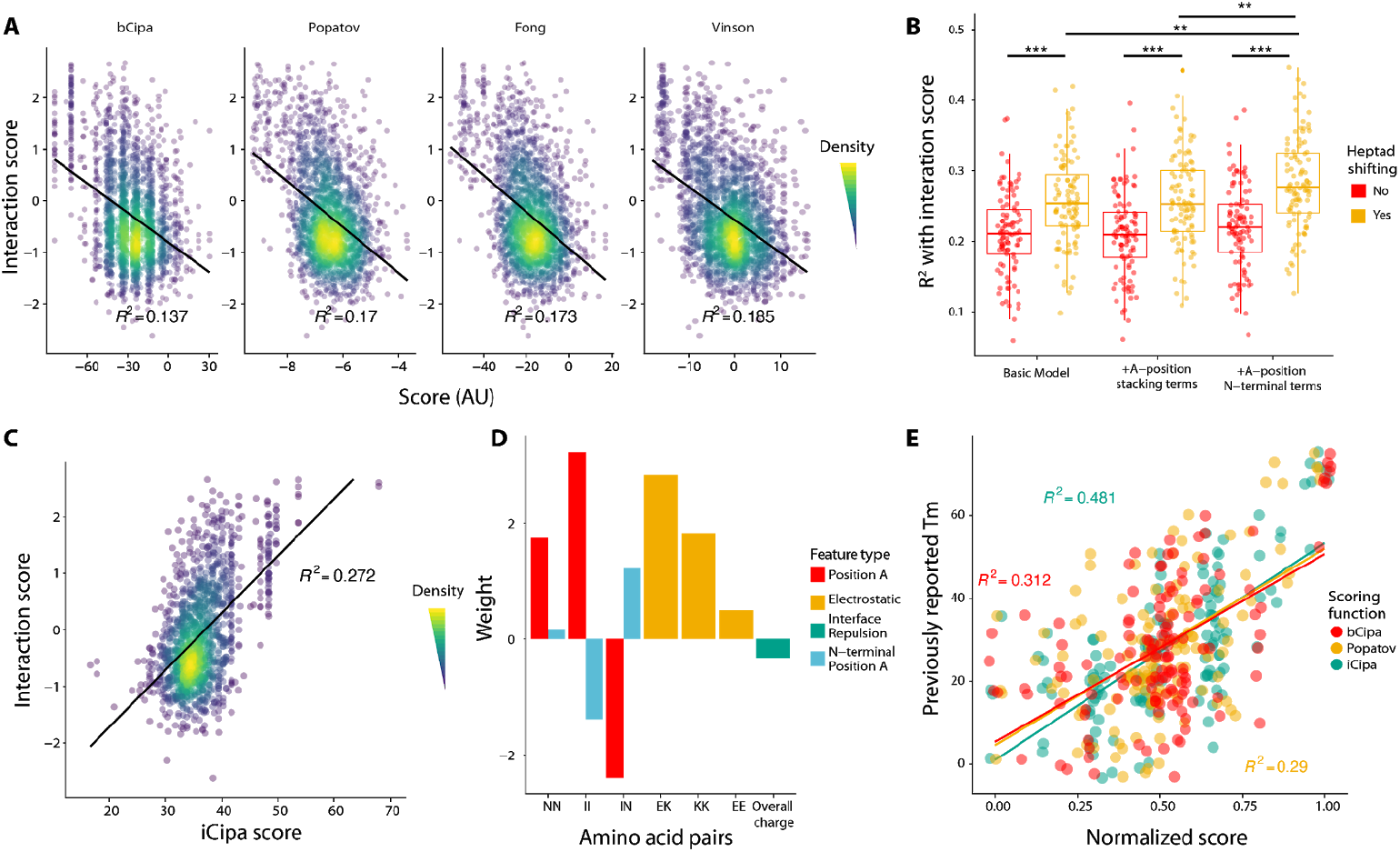
Comparison, development and validation of iCipa model. A) Previous models of coiled-coil interactions are only mildly predictive of protein interactions. All R2’s are Interaction Score as predicted by the scoring algorithms. The black line represents a linear model of Interaction scores predicted by the different algorithms. B) Coefficient of determination of Interaction scores with different iCipa candidates. Each point represents a subsample of ten percent of the total data. *** = p < 0.001, ** = p < 0.01, two-tailed t-test. C) iCipa is more predictive of Interaction score (R^2^ > 0.27) than previous models, shown in A. Black line represents a linear model of Interaction scores as predicted by iCipa scores D) Weights for the iCipa model. Each weight scores a single type pair of amino acid between the two interacting coiled-coil s. E) iCipa is more predictive of previously published CC0 Library melting points than the bCipa or Potapov algorithm. Individual dots represent previously reported melting points compared with the normalized score from one of three scoring algorithms.

iCipa is a linear model, which facilitates interpretation. The weights of iCipa have expected and unexpected characteristics (Figure 3D). a-position residues prefer Ile/Ile pairings and tolerate Asn/Asn pairings between proteins and disfavor Ile/Asn pairings as expected. As expected, e- and g-positions favor salt bridges between Glu/Lys and disfavor Glu/Glu pairings, but counterintuitively, Lys/Lys pairings are acceptable for forming the interface which may be due to non-canonical interactions by long side chain of lysine.

To test the iCipa model, we excluded all the data from the original CC0 Library while we trained the weights. When the scoring functions are normalized and compared (Figure 3E), both the Potapov/SVR and bCipa algorithms performed worse in predicting the measured melting points with R^2^ < 0.32 compared to iCipa, R^2^ = 0.48—a fifty percent increase in predictive ability. Importantly, the increase in predictive power for iCipa on the CC0 Library demonstrates that iCipa has not been trained on an artifact of the NGB2H system but that the NGB2H system provides high quality data on PPIs which can provide general insights into coiled-coil function.

### CCmax Library design and verification

To evaluate iCipa’s prediction capabilities, demonstrate the scalability of the NGB2H system, and identify larger orthogonal sets of coiled-coils, we built another library, the CCmax Library. The CCmax Library contains 18,491 interactions of 931 different coiled-coils in fifteen predicted orthogonal sets and seven control sets (Figure 4A). The orthogonal sets were designed using our computational framework and scored with one of fifteen variants of iCipa. After designing (Figure S2D) and cloning (Figure S3C, S4) the CCmax library, we collected high quality data on 17,320 interactions (Figure S15). The CC0 Library was a subset of the CCmax Library and it broadly agreed with its performance in our previous libraries (Figure S16).

**Figure 4).**
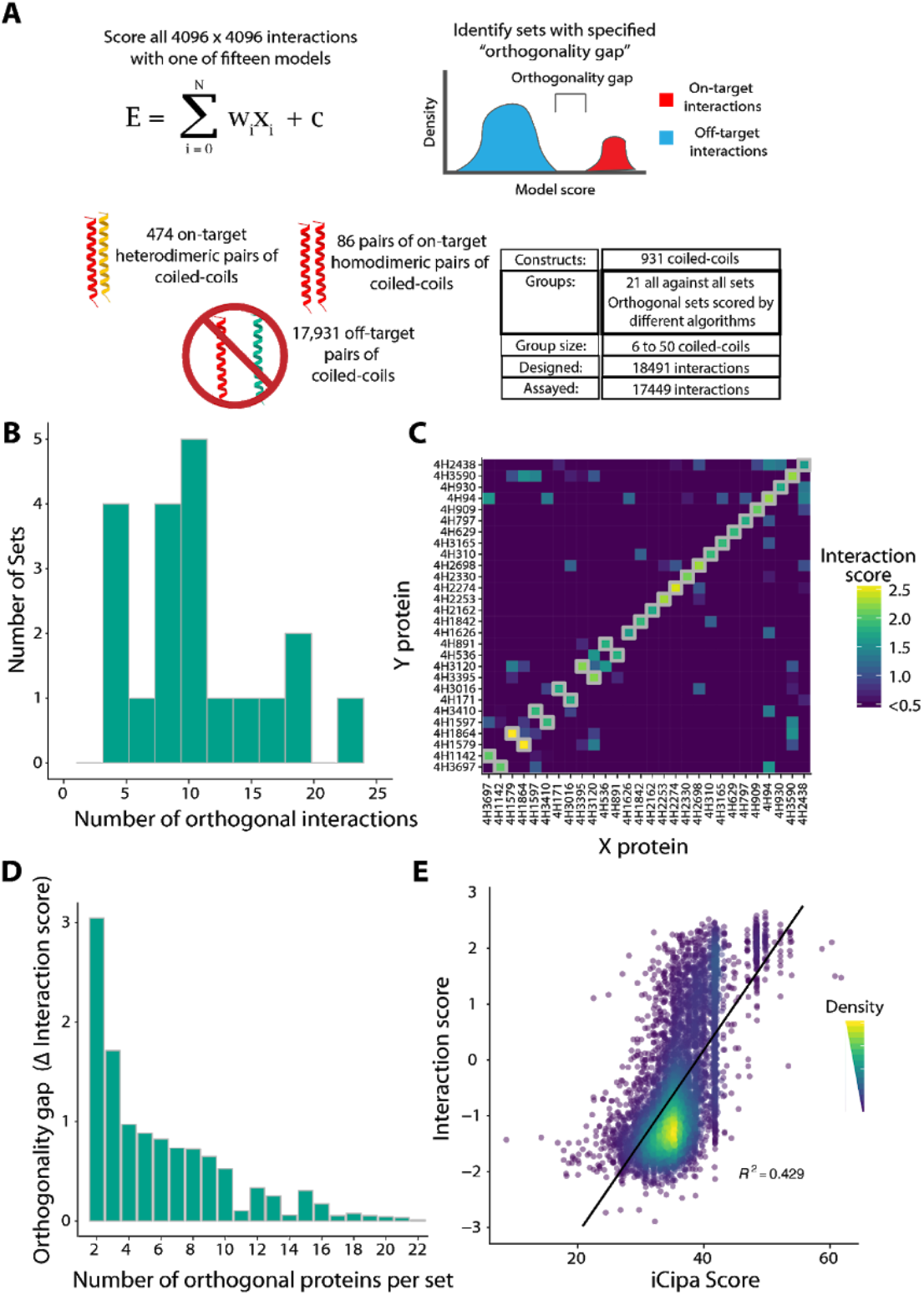
The largest orthogonal sets of the CCmax Library. A) Design of the CCmax library. Using seven different algorithms, each possible interaction was scored and sets of orthogonal interactions with an orthogonality gap were identified. In total 21 sets comprised of 18,491 interactions were analyzed. B) Number of on-target orthogonal interactions per set. Between 3 and 22 orthogonal interactions were obtained per set. C) The largest orthogonal set contains 22 on-target interactions across 784 tested pairs. Grey boxes represent on-target interactions. Un boxed squares represent non-interactions D) The largest orthogonality gap per number on-target interactions in a set. The orthogonality gap is the difference between the weakest on-target Interaction score and strongest off-target Interaction Score. E) iCipa’s agreement with the Interaction score (R2 = 0.429). The black line is a linear model predicting Interaction scores from iCipa predictions.

### Orthogonal sets of the CCmax Library

We identified orthogonal sets with sizes of up to twenty-two on-target pairs (Figure 4B) and 784 total interactions (Figure S17). Five of the sets contained more on-target interactions than any set in the CCNG1 Library, and fifteen contained more on-target interactions than the largest published set^15^. Our largest orthogonal set (Figure 4C) contained twenty-two coiled-coil dimers, sixteen homodimers and six heterodimers, which is ten on-target interactions larger than the state-of-the-art set from CCNG1. To characterize the accuracy of different iCipa variants, we subsampled each set (ten proteins, 500 times), and found the largest orthogonal set per subsampling. We found little significant difference between the algorithms (Figure S18) suggesting that orthogonality is still challenging to design using current algorithmic accuracy and underscoring the necessity of large scale experimental verification.

Different applications need varying levels of orthogonality; while gene circuits likely need extreme orthogonality, protein origami, which benefits from avidity, is not under such constraints. Thus, we identified the largest orthogonality gap for different numbers of on-target interactions. (Figure 4D, Table S7). As expected, smaller sets had larger gaps, but large orthogonality gaps were identified for sets as large as sixteen on-target interactions. Finally, we compared the CCmax Library’s Interaction score with iCipa predictions which show substantial improvement over the CCNG1 Library. iCipa was able to predict Interaction scores with R^2^ = 0.43 (Figure 4E). We attribute the increase in iCipa’s power to the use of a single coiled-coil backbone which consists of only alanine residues at the b-, c- and f-positions. The improvement in predictive power appeared in other algorithms to a lesser extent, all of which maintained an R^2^ < 0.28 (Figure S19).

## Discussion

We have developed and validated a novel system for high-throughput identification of PPIs. We built a framework to predict orthogonal coiled-coil interactions and used it to design tens of thousands of interactions which we then assayed with the NGB2H system in a design-build-test cycle. Using the data collected, we improved state-of-the-art coiled-coil interaction prediction algorithms which allowed us to design the largest set of any orthogonal proteins to date with twenty-two on-target interactions. Thus, by iterative design we demonstrate how high-throughput PPI identification can facilitate identification of desired protein function and improve design.

Our work builds on previous high-throughput two-hybrids to create a generalizable system for studying PPIs, that could include both soluble and membrane proteins. By uniting gene synthesis with a mapping step and a barcode readout, our system allows high throughput characterization of any binary PPI. Previous high-throughput studies used highly constrained libraries--either the ORFome^21–24^ of one of a handful of reference genomes, targeted single residue mutations which only explore a sliver of sequence space around a primary sequence^25,26^ or several randomly sheared coding sequences^27^. Using the capabilities of DNA synthesis broadens the testable sequence space which facilitates investigations of a variety of areas such as protein domains, extant genetic variation, evolutionary trajectories or epistatic effects. Furthermore, for the investigator who is not interested in an all-against-all approach, synthesis allows the explicit pairings of only certain proteins. While we benefited from the short length of our proteins of interest, recent pooled gene synthesis techniques^28,29^ can be used to interrogate much larger proteins. Deconvoluting library diversity has also been a challenge for other multiplexed assays. Other multiplexed methods involved picking colonies and sanger sequencing them^21^, mapping the beginning of reading frames to reference genomes^22–24^ or manually BLASTing obtained reads^27^. Our explicit mapping step allows for the high-throughput creation of a library to map arbitrary proteins to DNA barcodes, and because it is a separate step it could use long read sequencing to overcome the length limitations of Illumina sequencing. Finally, by using a barcode readout downstream of synthesis and mapping we can measure protein libraries in many formats.

Our improvements to coiled-coil design algorithms represent an important advance for *de novo* protein design. Though coiled-coil interactions have been modelled with diverse approaches, our iCipa algorithm shows clear advantages over existing models. In particular, heptad shifting provides an intuitive, biologically rational addition that can be applied to any future improvements in coiled-coil design. Overall, we found iCipa to be substantially more accurate than other tested algorithms, at least for this limited set of residues tested.

Here, we simultaneously performed a massive characterization of PPIs within a protein family and identified the largest set of orthogonal proteins identified to date. The CCmax Library characterized three times as many interactions than any previous intra-protein family work^13^. From the total of 26,049 interactions we characterized, we found a large number of orthogonal proteins—in sets of up to 12 heterodimers or 22 heterodimers and homodimers. Though orthogonal coiled-coils are particularly needed as the building blocks for protein origami^4,5^, they could be substituted for histidine kinases in orthogonal signaling pathways^30^, synthetic orthogonal transcriptional logic gates^8,31,32^, or for orthogonal cellular localization^33^.

Thus, the ability to characterize constructs across highly-diverse sequence space and for the identification of networked properties such as orthogonality, highlights the NGB2H’s scalability and generality. Because it can be adapted to any sequence the experimenter desires, the NGB2H facilitates interrogation of PPIs beyond endogenous interactomes, it can be used to characterize whole protein families, empirically inform protein design, or investigate complex phenomena like epistasis.

## Acknowledgements

We thank the members of the Kosuri and Plesa labs for their feedback on the manuscript and figures. We thank Suhua Feng of the UCLA Broad Stem Cell Research Center and the team of the Technology Center for Genomics and Bioinformatics for performing next-generation sequencing. We thank Octant Inc., the Kruglyak Lab at UCLA, and the Black Lab at UCLA, for use of their next-generation sequencers. We thank Mathew Graf and Will Silkworth for their assistance at the UCLA-DOE Biochemistry Shared Instrumentation Facility. We thank Thomas Kuhlman for kindly providing strain TK310. Finally, we thank Chris Voigt for sharing repressor/promoter sequences with us.

## Author contributions

H.K., W.C.B., N.L. and S.K. designed the NGB2H system. A.L., D.S. and R.J. designed the large sets of coiled-coils. H.K., N.L. and W.C.B. designed the oligonucleotide libraries. H.K. W.C.B. and J.L. performed the experiments. A.L. designed the improved interaction algorithms. W.C.B. and N.L. performed the computational analysis. W.C.B, A.L., R.J. and S.K. analyzed the results and iteratively planned the next steps. W.C.B created the figures. S.K., W.C.B. and A.L. wrote the manuscript with input from all authors.

## Methods

### Oligonucleotide designs

Libraries were designed as shown in Figure S2. Though the CC0 and CC1 Libraries were assembled from two oligonucleotides and the CCNG1 and CCmax Libraries from one oligonucleotide, they followed the same overall assembly logic. In brief, each library was flanked with two orthogonal 15bp primers^34^ for amplification from the OLS pool. Interior to the flanking primers were type IIS restriction enzyme sites to facilitate scarless cloning, and the complete coiled-coil sequence. The CC0 and CC1 Libraries contain extra type IIS sites and flanking 15bp primers to allow linking and amplification of the X and Y halves of the two-hybrid. A complete description of each design is listed in Supplementary Information Section 2 and all oligonucleotides used are listed in Table S1 while all proteins used are listed in Table S2.

### Orthogonal coiled-coil interaction prediction

To predict orthogonal coiled-coils, we generated all 4,096 possible four heptad coiled-coil with asparagine or isoleucine at the a-position and glutamic acid or lysine at the e- and g-positions and scored 16.7 million interactions in an all-on-all design using the Potapov algorithm (CCNG1 Library) or our iCipa candidate algorithms (CCmax Library). Calculating orthogonality is a challenging problem that scales in exponential time with the number of possible binding partners. We used a maximal clique algorithm to identify sets of orthogonal coiled-coils where all on-target interactions have a higher score than all off-target interactions and it runs in dozens seconds on a standard laptop. Full code can be found at: https://github.com/dancsi/DiplomaThesis

### Construct and library cloning

Each library was cloned in a similar manner, with slight differences in methods to attach a random DNA barcode to the OLS pools. After the 20bp of random DNA was attached with PCR to the 3’ end of the X and Y construct (Figure S3), constructs were sequenced in bulk on a MiSeq to identify it and a specific X and Y (below). After barcode mapping, the T25 and T18 + GFP halves were cloned in sequentially with type IIS restriction enzymes for scarless cloning (Figure S4). All enzymes and polymerases came from NEB. A complete description for how each library was cloned can be found in the Supplementary Information Section 4 and oligonucleotides used for cloning are listed in Table S3.

### Mapping random barcodes

Once random barcodes were attached and cloned, constructs were sequenced on an Illumina MiSeq to identify the X and Y proteins which each barcode was connected to. DNA containing the X and Y proteins, and the barcode were amplified as a linear fragment, and Illumina’s P5 and P7 adapters attached. Constructs were sequenced with a v3 300 cycle paired end kit (Illumina TG-142-3003), with custom primers spiked into the Illumina primers. Sequences were demultiplexed, and mapped with a BBtools pipeline and consensus building custom script. Full descriptions of how each library was mapped can be found in the Supplementary Information Section 5 and scripts can be found at https://github.com/cliff-b/ortho-ccs.

### Strains used

All NGB2H experiments were run in TK310^35^ carrying pSK34. TK310 is a previously published MG1655 derivative with deletions in cpdA, lacY and cyaA, which give it a large linear response range to cAMP. pSK34 contains repressors for both the phlF and Tet promoters to maintain repression of the two-hybrid proteins. CB216 is a NEB5ɑ derivative with pSK34 integrated genomically and only used for cloning. All plasmids used for basic cloning are listed in Table S4 and available at Addgene.

### NGB2H assay execution

Glycerol stocks of each library were thawed, and 100uL were grown up overnight in 100mL MOPS EZ Rich Defined Media (Teknova M2105) with kanamycin (Teknova K2125) and carbenicillin (Teknova C2130). For time course studies, a glycerol stock containing a library of constitutive GFP constructs was also thawed, and 100uL was inoculated into 10mL of MOPS EZ Rich Defined Media with kanamycin and carbenicillin and grown overnight. The next morning 1mL of the GFP library was added to the 100mL of library culture. After mixing GFP and experimental libraries, 1mL of overnight culture was added to a fresh culture of 100mL MOPS EZ Rich Defined Media with carbenicillin and kanamycin and the inducers for two hybrid expression: 5ng/mL anhydrotetracycline, 1.5uM 2,4-Diacylphlorolglucinol and 100uM IPTG, done twice for biological replicates, except where indicated (Supplemental Information Section 6). Flasks were placed in a 37C degree shaker for six hours. Samples were pulled after 6 hours and placed on an ice slurry to quickly cool for 15 minutes after which cells were spun down for RNA and DNA extraction.

### RNA and DNA preparation for barcode sequencing

Samples of RNA were prepared with Qiagen RNeasy kits (Qiagen 74106, or 75144) according to manufacturer’s instructions, with on column DNase digestion (Qiagen 79254) and concentrated with RNeasy MinElute Cleanup kit (Qiagen 74204). RNA was reverse transcribed with Superscript IV (ThermoFisher 18090050) with a modified protocol such that 25ug of input RNA was used, the extension step ran for 1h at 55C, and 1uL of RNase A was added in the RNA removal step. Each sample was transcribed with a specific primer, often oSK193 or oSK194, that attached the i7 index and P7 sequencing primer. Samples of DNA were prepared with Qiagen Plasmid Plus Maxi kits (Qiagen 12963) according to the manufacturer’s instructions. RNA samples were verified to contain very low levels of DNA (< 1:1000) by qPCR (Kapa Biotechnology KK4601) with oSK199 and oSK200, which was repeated with a high-fidelity PCR for a low number of cycles to keep samples in the exponential amplification phase. DNA samples were similarly quantified with qPCR and amplified for low cycles to attach P5 and P7 and multiplexing indices. Amplified samples were then quantified on an Agilent Tapestation 2200 with D1000 screentape (Agilent 5067-5582), verified to be monodispersed and mixed in equimolar quantities. Complete details for RNA and DNA preparation can be found in the Supplementary Information Section 7.

### Barcode sequencing

Pooled RNA and DNA barcodes from each experiment were sequenced with various cores and startups at UCLA. The CC1 and CCNG1 Libraries were sequenced on a Hiseq 2500 while the CCmax and CC0 libraries were sequenced on a Nextseq 550. Samples were diluted and mixed with 5-20% phiX control v3 (Illumina FC-110-3001) and sequenced with oSK326 for read 1 and oSK324 for the index read.

### Barcode Counting

We used a custom bash script to count DNA barcodes from barcode sequencing. After demultiplexing into reads from RNA or DNA samples, reads were truncated to the 20bp containing the barcode and unique sequences counted. Barcode counts were then processed with Starcode (v1.3), to condense barcodes within a levenshtein distance of one to remove sequencing errors and tallied again. Full processing scripts can be found at https://github.com/cliff-b/ortho-ccs.

### Interaction quantification

Barcode count files were imported into R where they were merged with the mapping file to provide the protein pair identified with each barcode. Barcodes corresponding to the same construct were summarized (dplyr 0.7.4) and total counts of RNA barcodes and DNA barcodes per protein pair were obtained. For our analysis we used Interactions scores calculated as 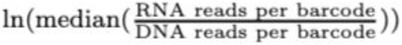 for barcodes that had >10 reads in all DNA samples. Interactions for all libraries are reported in Table S5.

### Orthogonal set identification

Orthogonal sets were identified for the CCNG1 and CCmax libraries. Briefly, we wrote a script, maximal.py that took the Interaction scores for each set and built a graph with interactions forming the edges between proteins. Finding the maximum independent set of the line graph of this graph gave us the largest orthogonal set of interactions, listed in Table S6. The largest sets for different numbers of on-target interactions are listed in Table S7. The full script for orthogonal set identification can be found at https://github.com/cliff-b/ortho-ccs.

## References

1. Vidal, M., Cusick, M. E. & Barabási, A.-L. Interactome networks and human disease. Cell 144, 986–98 (2011).

2. Huang, P.-S., Boyken, S. E. & Baker, D. The coming of age of de novo protein design. Nature 537, 320–7 (2016).

3. Ljubetič, A., Gradišar, H. & Jerala, R. Advances in design of protein folds and assemblies. Curr. Opin. Chem. Biol. 40, 65–71 (2017).

4. Ljubetič, A. et al. Design of coiled-coil protein-origami cages that self-assemble in vitro and in vivo. Nat. Biotechnol. 35, 1094–1101 (2017).

5. Gradišar, H. et al. Design of a single-chain polypeptide tetrahedron assembled from coiled-coil segments. Nat. Chem. Biol. 9, 362–6 (2013).

6. Boyken, S. E. et al. De novo design of protein homo-oligomers with modular hydrogen-bond network – mediated specificity. 399, 69–72 (2016).

7. Fallas, J. A. et al. Computational design of self-assembling cyclic protein homo-oligomers. Nat. Chem. 9, 353–360 (2017).

8. Chen, Z. et al. Programmable design of orthogonal protein heterodimers. Nature 565, 106–111 (2019).

9. Pauling, L. & Corey, R. B. COMPOUND HELICAL CONFIGURATIONS OF POLYPEPTIDE CHAINS: STRUCTURE OF PROTEINS OF THE ex-KERATIN TYPE. 3.

10. Crick, F. H. C. The packing of α-helices: simple coiled-coils. Acta Crystallogr. 6, 689–697 (1953).

11. Crick, F. H. C. The Fourier transform of a coiled-coil. Acta Crystallogr. 6, 685–689 (1953).

12. Acharya, A., Rishi, V. & Vinson, C. Stability of 100 homo and heterotypic coiled-coil a-a′ pairs for ten amino acids (A, L, I, V, N, K, S, T, E, and R). Biochemistry 45, 11324–11332 (2006).

13. Potapov, V., Kaplan, J. B. & Keating, A. E. Data-Driven Prediction and Design of bZIP Coiled-Coil Interactions. PLoS Comput. Biol. 11, 1–28 (2015).

14. Mason, J. M., Schmitz, M. a, Müller, K. M. & Arndt, K. M. Semirational design of Jun-Fos coiled coils with increased affinity: Universal implications for leucine zipper prediction and design. Proc. Natl. Acad. Sci. U. S. A. 103, 8989–8994 (2006).

15. Crooks, R. O., Lathbridge, A., Panek, A. S. & Mason, J. M. Computational Prediction and Design for Creating Iteratively Larger Heterospecific Coiled Coil Sets. Biochemistry 56, 1573–1584 (2017).

16. Thompson, K. E., Bashor, C. J., Lim, W. A. & Keating, A. E. SYNZIP Protein Interaction Toolbox: in Vitro and in Vivo Specifications of Heterospecific Coiled-Coil Interaction Domains. ACS Synth. Biol. 1, 118–129 (2012).

17. Karimova, G., Pidoux, J., Ullmann, a & Ladant, D. A bacterial two-hybrid system based on a reconstituted signal transduction pathway. Proc. Natl. Acad. Sci. U. S. A. 95, 5752–5756 (1998).

18. Brodnik, A., Jovičić, V., Palangetić, M. & Silađi, D. Construction of orthogonal CC-sets. Informatica 43, (2019).

19. Drobnak, I., Gradišar, H., Ljubetič, A., Merljak, E. & Jerala, R. Modulation of Coiled-Coil Dimer Stability through Surface Residues while Preserving Pairing Specificity. J. Am. Chem. Soc. 139, 8229–8236 (2017).

20. Fong, J., Keating, A. & Singh, M. Predicting specificity in bZIP coiled-coil protein interactions. Genome Biol. 5, R11 (2004).

21. Yachie, N. et al. Pooled-matrix protein interaction screens using Barcode Fusion Genetics. Mol. Syst. Biol. 12, 863–863 (2016).

22. Trigg, S. A. et al. CrY2H-seq: A massively multiplexed assay for deep-coverage interactome mapping. Nat. Methods 14, 819–825 (2017).

23. Yang, J.-S. et al. rec-YnH enables simultaneous many-by-many detection of direct protein–protein and protein–RNA interactions. Nat. Commun. 9, 3747 (2018).

24. Yang, F. et al. Development and application of a recombination-based library versus library high-throughput yeast two-hybrid (RLL-Y2H) screening system. Nucleic Acids Res. 46, e17–e17 (2018).

25. Younger, D., Berger, S., Baker, D. & Klavins, E. High-throughput characterization of protein–protein interactions by reprogramming yeast mating. Proc. Natl. Acad. Sci. 114, 12166–12171 (2017).

26. Diss, G. & Lehner, B. The genetic landscape of a physical interaction. eLife 7, 1–31 (2018).

27. Andrews, S. S. et al. High-resolution protein–protein interaction mapping using all- *versus* -all sequencing (AVA-Seq). J. Biol. Chem. 294, 11549–11558 (2019).

28. Plesa, C., Sidore, A. M., Lubock, N. B., Zhang, D. & Kosuri, S. Multiplexed gene synthesis in emulsions for exploring protein functional landscapes. 6 (2018).

29. Sidore, A. M., Plesa, C., Samson, J. A., Lubock, N. B. & Kosuri, S. DropSynth 2.0: high-fidelity multiplexed gene synthesis in emulsions. Nucleic Acids Res. 48, e95–e95 (2020).

30. McClune, C. J., Alvarez-Buylla, A., Voigt, C. A. & Laub, M. T. Engineering orthogonal signalling pathways reveals the sparse occupancy of sequence space. Nature 574, 702–706 (2019).

31. Chen, Z. et al. De novo design of protein logic gates. Science 368, 78–84 (2020).

32. Fink, T. et al. Design of fast proteolysis-based signaling and logic circuits in mammalian cells. Nat. Chem. Biol. 15, 115–122 (2019).

33. Lebar, T., Lainšček, D., Merljak, E., Aupič, J. & Jerala, R. A tunable orthogonal coiled-coil interaction toolbox for engineering mammalian cells. Nat. Chem. Biol. 16, 513–519 (2020).

34. Kosuri, S. et al. Scalable gene synthesis by selective amplification of DNA pools from high-fidelity microchips. Nat. Biotechnol. 28, 1295–1299 (2010).

35. Kuhlman, T., Zhang, Z., Saier, M. H. & Hwa, T. Combinatorial transcriptional control of the lactose operon of Escherichia coli. Proc. Natl. Acad. Sci. U. S. A. 104, 6043–8 (2007).

