## Supplemental Information for "A Multiplexed Bacterial Two-Hybrid for Rapid Characterization of Protein-Protein Interactions and Iterative Protein Design"

Supplementary Information

### 1. System design

#### 1.1 NGB2H system design

We decided to use the *B. pertussis* adenylate cyclase bacterial two-hybrid[1], as the starting point for our high-throughput two-hybrid to access the genetic tools available in *E. coli*. To make a quantitative, inducible, multiplexed system we made numerous changes to Karimova et al.'s two-hybrid. First, the two halves of adenylate cyclase, T18 and T25, are traditionally expressed on separate plasmids, which is incompatible with pooled library screens in *E. coli*. Second, both T18 and T25 were under control of a lacUV5 promoter which creates a positive feedback loop because it is cAMP inducible[2]. Though the authors note that a thresholding effect is useful for their work, this prohibits a quantitative PPI screen. Third, the presence of exogenous cAMP confers a growth advantage except during transformation when it is toxic (unpublished data) which could lead to highly biased libraries. To simultaneously maintain plasmid stability with both halves of adenylate cyclase on the same plasmid, remove the positive feedback loop, and ameliorate cAMP growth effects, and, we placed T18 and T25 under separate inducible promoters, pPhlF and pTet[3], respectively. Though pPhlF and pTet did not originally have the same induced expression levels, by tuning their ribosome binding sites we were able to equalize their expression. Finally, as T18 and T25 were on the same plasmid as the reporter, we were concerned about efficient termination, so all transcripts ended with two strong terminators[4].

In addition to our modifications to adenylate cyclase we built a new reporter to create a multiplexed assay with the genetic elements necessary for an Illumina-based readout. Most significantly, we replaced LacZ with sfGFP to allow efficient measurements in a standard plate reader of single interactions. This allowed for us to characterize kinetics of the NGB2H system, quickly optimize genetic elements and test single constructs. We also changed the promoter to a variant of pLac[5] which had the greatest fold induction (Figure S1B, S1C), to increase the system's dynamic range. To enable Illumina sequencing of reporter expression we placed a Truseq adaptor, a placeholder for a DNA barcode, and reverse transcription primers in the 3' untranslated region of sfGFP. In addition to our episomal changes, we replaced the original strains with TK310, an MG1655 derivative with deletions in adenylate cyclase, lactose permease and cAMP-phosphodiesterase which gives the Lac promoter a linear response to cAMP[6], enabling direct comparison between interactions. Finally, for studies in TK310 we included a plasmid with the PhlF and Tet repressors, which we call pSK34.

### 2. Library composition and design

#### 2.1 CC0 Library design

Jody Mason and colleagues used circular dichroism to characterize a set 16x16 of orthogonal coiled-coils[7] which we used for validation of the NGB2H system. To create the CC0 library, which consisted of all 256 interactions, we used a custom python script difflength-lib-gen.py with mason.fasta to specify the protein sequences. This script created 230bp oligonucleotides encoding either the X protein or Y protein (Figure S2A). We noticed that the chip amplification of some proteins would fail for unknown reasons with certain primers so we designed a redundant system of five subpools containing the same protein encoding DNA. The X containing oligonucleotides were designed to contain one of five sets of flanking subpool primers: oSK528-oSK532 for the forward primers and oSK609-oSK613 for the reverse primers.

Immediately 3' to the forward subpool primer was a group forward subpool primer, oSK608 which was the same across subpools. Downstream of the primers was the reverse complement of one of three codon usages for each CC0 coiled-coil, followed by a BspQI site for scarless cloning into the NGB2H system. Finally, the BspQI site was followed by a BbsI site that would allow ligation with oligonucleotides containing the Y protein and a spacer to make the oligonucleotide 230bp.

The oligonucleotides containing the Y protein were similarly designed. The Y containing oligonucleotides were flanked with one of five sets of subpool primers oSK533-oSK537 for the forward primer and oSK614-oSK618 for the reverse primers. Immediately 3' to the forward primer is a spacer to make the oligonucleotides 230bp, followed by a BbsI site to allow ligation with the X containing oligonucleotides. Downstream of the BbsI site was a Bts $\alpha$ I site to allow scarless cloning of the Y oligonucleotides into the NGB2H system followed by one of three codon usages for the CC0 coiled-coils. At the end of the coiled-coil protein coding sequence we placed a BsaI site to allow insertion of the T18 half of the *B. pertussis* two-hybrid. Finally, 5' adjacent to the reverse subpool primer, we placed a group subpool reverse primer oSK687. The five subpools with three codon usages of sixteen proteins in two orientations for a total of 480 oligonucleotides were ordered as an OLS pool from Agilent's high-fidelity process.

##### 2.2 CC1 Library composition

We designed a library of twenty coiled-coils based on our own designs, which we call the CC1 Library (Figure S9A). Half of the CC1 Library, P# set, has previously been partially characterized[8]. We expected the P# set to be orthogonal based on Glu/Lys bonding from the E- and G-positions and Ile/Ile or Asn/Asn complementarity at the A-position. The other half of the CC1 Library are backbone variants (changes at the B, C, and F-positions) of four and six coiled-coils, the P#SC# set, and P#mS set respectively. As the B, C, and F-positions are thought to contribute little to coiled-coil specificity, we expected the P#mS and P#SC# constructs to behave like the corresponding P# constructs.

##### 2.3 CC1 Library design

The proteins were converted into oligonucleotides for either the X or Y half of the two-hybrid, with the accessory sequences for cloning into the NGB2H system (Figure S2B) with a custom script, codon_opt_hb.py. Briefly, the protein sequences were reverse translated into codon optimized DNA sequences and the sequence was checked for restriction enzyme sites used in downstream cloning steps and replaced any found.

We composed the X oligonucleotides from 5' to 3' thusly: a forward OLS amplification primer, stop codon, the reverse complement of the protein coding sequence, a BspQI site for scarless cloning, a junk spacer region, a reverse OLS amplification primer and another spacer to extend the oligonucleotides to a uniform 230bp. We used different amplification primers for different codon usages: oSK529 with oSK699, oSK700 with oSK701 or oSK702 with oSK703.

We composed the Y oligonucleotides from 5' to 3' thusly: a forward OLS amplification primer, a junk spacer region, a Bts $\alpha$ I site to facilitate scarless cloning, the coding sequence of the protein, a BsaI site for scarless cloning, a reverse OLS amplification primer and a spacer base to make the oligonucleotides a uniform 230bp. We used different amplification primers for different codon usages: oSK570 with oSK704, oSK705 with oSK706 and oSK708 with oSK709. The twenty one proteins, with three codon usages in two orientations were ordered as 126 oligonucleotides from Agilent as an OLS pool.

#### 2.4 CCNG1 Library composition

The CCNG1 Library was composed of four heptad coiled-coils with either one or two Asn in the A-position with the other A-positions being Ile. E- and G-positions varied between Glu and Lys with no limits on abundance. All D-positions were Leu to improve dimerization. With these constraints, our orthogonal computational framework (below) identified fifteen potential orthogonal sets to which we added previously tested designs, the P# set from the CC1 Library and the CC0 Library. We included six different backbone compositions (Figure S13A)--three across the experimental sets, two more in both control sets, and one in just the CC0 Library . These backbones were intended to vary stability and helical propensity: bA-largely alanine, bN-largely glutamine and bH-glutamic acid and lysine, bS-largely serine and glutamine and bP-largely lysine and glutamic acid, as well as the published backbone on the CC0 Library. The CCNG1 Library encoded 132 on-target homodimers, 429 on-target heterodimers and 7,608 off-target interactions.

#### 2.5 CCNG1 Library design

To create the CCNG1 Library we used a custom python script, lib-gen.py which produced 230bp oligonucleotides containing each pair of proteins specified in 170307 all-8k R8000.pairs with the protein coding sequence in 170307 all-8k R8000.fasta. This script created an oligonucleotide that started with oSK229 followed by the reverse complement of one of three codon usages of the X protein (Figure S2C). Downstream of the X protein was a BspQI site to allow scarless cloning of the X protein and a Bts$\alpha$I site to allow scarless cloning of the Y protein. The Y protein followed the Bts$\alpha$I site in one of three codon usages before terminating in a BsaI site to allow scarless cloning of the T18 half of the NGB2H assay. Finally, at the 3' end, it contained the reverse complement of oSK232. The script checked each protein coding region for restriction enzyme sites, remaking those that contained a reserved enzyme site, and added one nucleotide to bring the constructs to a uniform 230bp. The 24,483 oligonucleotides were ordered from Agilent as an OLS pool.

#### 2.6 CCmax Library composition

The CCmax Library was largely composed under the same constraints as the CCNG1 Library. Library members were four heptad coiled-coils with the A-positions varying between Ile and Asn, the D-position invariantly Leu and the E- and G-positions varying between Glu and Lys. The backbones were invariantly Alanine. We also included two sets of anti-parallel coiled-coils under the analogous constraints. Our computational pipeline used fifteen different iCipa candidates as scoring functions and we took the largest orthogonal set produced by each. We also included five control sets that had been previously evaluated, including the CC0 Library and P# set from the CC1 Library. It also included two large sets that systematically varied each A-, E- and G- position though but were not predicted to be orthogonal and thus excluded from the analysis. The CCmax Library encoded 474 on-target heterodimers, 86 pairs of on-target homodimers and 17,931 off-target interactions for a total of 18,491 interactions.

2.7 CCmax Library oligo design

To create the CCmax Library we used a custom python script, lib-gen2.py, which functioned very similarly to lib-gen.py. Briefly, it took all pairs of proteins in 18491_flipped_zipped_final_pairs.pairs and encoded them into DNA from proteins sequences in 18491_flipped_zipped_final_pairs.fasta into 230bp oligonucleotides with the functional DNA sequences needed for cloning into the NGB2H assay (Figure S2D). This created oligonucleotides flanked with oSK470 and oSK471 for subpool amplification. Immediately 5’ to oSK470 was the reverse complement of the X protein in one of three codon usages, all of which were optimized to the frequency which they naturally occur in E. coli. To facilitate scarless cloning the X protein was followed by a BspQI site. Likewise, for scarless cloning of the Y protein we placed a Bts $\alpha$ I site immediately downstream of the BspQI site and N-terminal to Y protein coding sequence. Finally, we added a BsaI site C-terminal to the Y protein coding sequence. We repeated this process three times to create three different codon usages and then ordered the full set of 55,473 oligonucleotides as an OLS pool from Agilent using their high-fidelity process.

2.8 GFP Library design

The GFP Library was designed as a library of barcodes that remain constant despite difference in library composition, similar to ERCC for RNA-seq[9]. We created a library of constitutive GFP constructs spanning several orders of magnitude in expression, that contained a unique DNA barcode and the flanking sequences for amplification and sequencing with our other libraries. We used previously published expression data[10] from which we selected constitutive promoters that would give a wide range in expression. In order from low to high expression we chose J23117, pAPFAB277, pAPFAB69, pAPFAB48, pAPFAB70 and pAPFAB101. Each of these promoters was attached to sfGFP and in the 3’ UTR of the sfGFP we inserted our Truseq adaptor and a 7bp random barcode.

#### 3. Computational framework and model refinement

3.1 Orthogonal computational framework

We built a computational framework to calculate predicted interactions for four heptad coiled-coils, for which all programs can be found at https://github.com/dancsi/DiplomaThesis. To limit the search space, we examined a subset of four heptad coiled coils where the A-positions varied between Ile and Asn and the E- and G-positions varied between Lys and Glu, for a total of 4096 possible sequences (2*2*2)^4. Using the algorithm described in Potapov et al.,[11] we scored all 16.7 million potential interactions. To make this computationally tractable we reimplemented the algorithm in the C for a 1000x fold increase in speed in fastscore.exe. After scoring interactions we sought to identify sets of orthogonal interactions, those where all on-target interactions had higher interaction scores than all off-target interactions. The identification of orthogonal interactions can be reduced to the maximum clique problem[12] which we implemented in Solver.exe. Solver.exe uses score cutoffs for on-target and off-target interactions to enforce orthogonality. Based on experimental data of the P# and CC0 sets we searched for sets with a difference of 1.0 (arbitrary units) between the scoring cutoffs. Finally, to obtain the multiple sets of the CCNG1 Library we varied the threshold for non-interacting pairs from -7.5 to -9 in increments of 0.05.

3.2 Model refinement from CCNG1 Library

We used the CC0 Library subset in the CCNG1 Library as a calibration curve to convert interaction scores to Tms to be comparable to other coiled-coil interaction prediction algorithms (Fig S11B). We found that the prediction power of the algorithm increases significantly if we allow coiled-coils to bind in the heptad alignment that maximizes the $\Delta$G. In calculating the off-target pairs, most algorithms assume the coiled-coil interacts with all four heptads but this does not necessarily reflect reality as a three heptad alignments may be more energetically favorable (Figure S14). To develop an algorithm that incorporates heptad shifts, we first extracted the features of all possible interactions without permitting shifts. Using this model we have scored three heptad shifts at positions -7, 0, and +7. We then changed the alignment to the lowest scoring position and retrained the model. After five iterations the model parameters converged on a fixed point.

We fit several groups of parameters in a series of models. First we incorporated features known to be important to coiled-coil interaction by counting the number of aligned residues between pairs of coiled-coils that were Asn/Asn, Asn/Ile, or Ile/Ile at the A-position, and Glu/Glu, Glu/Lys, or Lys/Lys between E- and G-positions. We also included a term for total charge between the two coiled-coils. As heptad shifting means fewer Leu in the D-position are interacting, we also scored all parameter groups with and without a term for the D-position. We then tested two separate parameter groups, one that looked at the effect of consecutive A-positions and one that looked at the most N-terminal A-position. Our consecutive A-position parameter group compared Ile/Ile, Asn/Asn, Asn/Ile and Ile/Asn in consecutive heptads, while our N-terminal parameter group looked at Ile/Ile, Asn/Asn and Asn/Ile pairs at the first heptad. Of note, the first heptad was scored again in the central suite. All models were fit using the Ridge regression algorithm of the sklearn library. Interactions were weighted by the number of different barcode counts, and on-target pairs upweighted 10 fold. We found that our term for repulsion to be beneficial to our core model, but scoring Leu interactions did not matter. Our consecutive A-position parameter group did not improve the core model's predictive capabilities but the N-terminal group significantly did (Figure 3B). Thus our best model, which we call iCipa, is the core model with N-terminal A-position parameters (Figure 3D).

Our model comes with several limitations. In order to limit the design space, the model only uses Glu/Lys at the E- and G-positions and Asn/Ile at the A positions while the D-position is expected to be a Leu and B-, C- and F-positions are expected to be Ala. As the backbone is expected to be Ala, the helical propensity of all proteins is very expected to be very high. Because of this, our model does not include a term for helical propensity as it was not predictive of interaction strength.

**4. Library Cloning**

For all cloning steps the reagents purchased from the following vendors unless otherwise noted. All reactions were performed according to the manufacturer's instructions unless otherwise noted. All restriction enzymes, phosphatase and ligase were purchased from NEB and DNA polymerase was NEBNext Q5 Hotstart HiFi PCR Master Mix (NEB M0543L) unless otherwise noted. All qPCR was performed using KAPA SYBR Fast 2x Master Mix (Kapa Biotechnology KK4601). All nucleic acid prep was purchased from Qiagen (Qiagen Plasmid Plus Maxi Kit 12963), though DNA cleanup and gel extraction kits were from Zymo (Zymo D4014 and Zymo D4008).

4.1 CC0 Library cloning

The 10pM high-fidelity OLS pool containing the lyophilized CC0 Library was resuspended in 25uL EB. This was diluted 1:20 in ddH2O and used as template for qPCR using the subpool primers oSK528-oSK532 and oSK609-oSK613 for X oligonucleotides and oSK533-oSK537 and oSK614-oSK618 for Y oligonucleotides. All subpools exhibited exponential amplification through 20 cycles, so high-fidelity PCR was performed in triplicate for 18 cycles. Replicates were pooled and digested at 1ug scale with BbsI-HF for two hours. Digested samples were run on a 4% agarose gel and the band containing the X or Y-protein was extracted. The entire extracted product was ligated with its subpool partner (ie X subpool-1 with Y subpool-1) with T4 ligase overnight at 50ng scale. Ligated samples were cleaned up and qPCR with the group pool primers, oSK608 and oSK687 established exponential amplification through eight cycles, with the exception of group pool one which showed no amplification and was excluded from downstream steps. High-fidelity PCR was performed for six cycles and cleaned up. Each group pool's concentration was analyzed with an Agilent Tapestation 2200 (Agilent 5067-5582) and equimolar fractions were pooled and diluted 100 fold. qPCR of the group pool samples with primers oSK689 to attach an AscI site and oSK690 to attach a BsaI site, random 20bp barcode, and EcoRI site, showed exponential amplification through ten cycles. High-fidelity PCR was performed in sextuplicat of which three were pooled to make the primary barcoding and three were pooled to make a replicate barcoding (Figure 1F). The barcoded products were run on a 3% agarose gel, extracted and digested at 2ug scale with AscI and EcoRI-HF for two hours. Freshly prepared pSK33 was likewise digested with AscI, EcoRI-HF and rSAP for two hours, run on a 1% agarose gel and extracted. Digested plasmid and barcoded product were ligated with T4 ligase for two hours at 250ng scale. Sample was cleaned up into 6uL ddH2O, and 1uL electroporated into NEB 5-alpha (NEB C2989K). After 35 minutes recovery in SOC samples were plated on LB agar + Kanamycin and grown overnight at 37C. Of the 2.4 million transformants, ~40,000 were scraped off the overnight plates and grown up in 150mL LB + Kanamycin at 30C overnight which was then purified and used for cloning the T25 segment.

The T25 insert was cloned as follows. The T25 segment was amplified from pSK59 with oSK694 and oSK695. PCR product was run on a 1% agarose gel and extracted. The insert was digested with BspQI for two hours at 4ug scale. Product was cleaned up and digested with BtsɑI for two hours. Plasmid from the above cloning step, for both barcodings, was digested with BspQI for three hours at 5ug scale, cleaned up and digested with BtsɑI overnight. Plasmid was then cooled to 37C and rSAP was added for 30 minutes. Dephosphorylated plasmid was run on a 1% agarose gel and extracted. The digested plasmids and T25 were ligated with T4 ligase at 250ng scale for six hours before transformation into freshly prepared electrocompetent CB216. Cells were recovered in SOC for 45 minutes and plated on LB agar + Kanamycin + Carbenicillin. The transformation was repeated three separate times for a total of ~100,000 colonies. These were scraped from the plates, diluted to OD 0.02 and grown up in 150mL LB + Kanamycin + Carbenicillin at 30C overnight which was then purified and used for cloning the T18 segment.

The T18 insert was cloned as follows. The T18 segment was amplified from pSK59 with oSK698 and oSK202. PCR product was run on a 1% agarose gel and a band corresponding to the expected 1715bp was extracted and digested with BsaI-HFv2 at 3ug scale for two hours. The vectors containing both barcodings were digested with BsaI-HFv2 and rSAP for two hours at 4ug scale. Digested plasmids were run on a 1% agarose gel and extracted. Plasmids and the T18 insert were ligated at 200ng scale with T4 ligase overnight. Ligation products were cleaned up and electroporated into freshly prepared electrocompetent CB216. Cells were recovered in SOC for 35 minutes and plated on LB agar + Kanamycin + Carbenicillin. Approximately 180,000 colonies were obtained for each barcoding. These were scraped from the plates diluted to OD 0.02 and grown up in 150mL LB + Kanamycin + Carbenicillin at 30C overnight. DNA was extracted, run on a 1% agarose gel, extracted, and electroporated at low concentration into previously frozen electrocompetent pSK34. Samples were recovered for 35 minutes in SOC at 37C and plated onto LB agar + Kanamycin + Carbencillin. Cells were scraped off the plates, diluted to OD 0.02 and grown up in 150mL LB + Kanamycin + Carbenicillin at 30C overnight. Glycerol stocks of overnight culture were prepared and stored at -80C. For downstream experiments one tube was fully thawed and subsequently discarded.

4.2 CC1 Library cloning

The 10pM OLS pool containing the CC1 Library was resuspended in 30uL EB. Each respective subpool—the X oligonucleotides with codon usages 1, 2 or 3 or the Y oligonucleotides with codon usages 1, 2, or 3—were amplified with their flanking primers with KAPA Real-time Library Amplification (KAPA Biosystems KK2702) via qPCR (oSK529 and oSK699, oSK700 and oSK701, oSK702 and oSK703 for X oligonucleotides and oSK570 and oSK704, oSK705 and oSK706, oSK708 and oSK709 for Y oligonucleotides). Using a 1:10 dilution of the OLS pool in ddH2O we found that the pools all exhibited exponential amplification through twenty-five cycles so amplification was repeated for twenty cycles. The products were cleaned up and qPCR was performed again with primers to attach a 30bp annealing region for the X oligonucleotides or a 30bp annealing region and the reverse primer from the first amplification for the Y oligonucleotides. This was done first with qPCR with KAPA Real-time Library Amplification which suggested exponential amplification through 10 cycles. It was then repeated with KAPA HiFI HotStart Ready for 10 cycles in quadruplicate, which were then purified and pooled. We then prepared a primerless PCR with mixtures of X and Y oligonucleotides of different codon usages. 100ng of X and Y were mixed into a 50uL KAPA HiFi HotStart Ready Mix reaction, with ten cycles of denaturation, annealing of the 30bp complementary regions, and extension. These samples were run on a 2.5% agarose gel and bands corresponding to the expected length of 400bp were extracted. Mixed samples were then amplified with biotinylated primers, oSK712, oSK713, or oSK714 and oSK715, oSK716 or oSK717 to attach restriction enzyme sites and barcodes using qPCR. Samples showed exponential amplification through fifteen cycles. The PCR was then repeated with KAPA HiFi HotStart Ready Mix for twelve cycles after which they were cleaned up and digested with AscI and EcoRI-HF at 0.4ug scale for two hours. Samples were heat inactivated and cleaned up with Dynabeads (ThermoFisher 65306) to remove undigested product. Likewise, backbone pSK33 was purified, PCR amplified with biotinylated primers oSK718 and oSK719 (KAPA Biosystems KK2602), digested with AscI and EcoRI-HF at 10ug scale for two hours and cleaned up with Dynabeads. The barcoded CC1 X and Y proteins were ligated into the digested vector with T4 Ligase at 100ng scale for an hour. Samples were cleaned up and electroporated into freshly prepared electrocompetent pSK34. After recovery in SOC media for an hour, cells were plated on LB agar + Kanamycin + Carbenicillin in serial 10-fold dilutions. The next day colonies were counted and the transformation was repeated while taking only 441,000 transformants (100x library coverage) from the SOC media. This was inoculated into 400mL of LB +Carbenicillin + Kanamycin at 30C overnight and DNA extracted for cloning the T25 insert.

The T25 insert was cloned into the barcoded X and Y proteins. It was amplified from pSK59 with biotinylated primers oSK203 and oSK204 with KAPA HiFi HotStart Ready Mix. 1.4ug was digested with BspQI and BtsαI in NEB Cutsmart buffer at 55C for one hour. Undigested product was removed with Dynabeads, and purified again. 30ug of barcoded plasmid product was digested with 10uL of BspQI and 10uL BtsαI in 200uL NEB Cutsmart for an hour. Product was run on a 1% agarose gel and the corresponding band extracted before dephosporylation with rSAP at 60uL scale for 30 minutes. 400ng of vector was ligated with T4 ligase for one hour, cleaned up, drop dialyzed and electroporated into pSK34 in two separate reactions. Colony PCR showed ~230k transformants, which were grown up in 200mL LB + Kanamycin + Carbenicillin overnight and plasmid was extracted for cloning in the T18 insert.

The T18 insert was amplified from pSK59 with biotinylated primers oSK201 and oSK202 with KAPA HiFi HotStart Ready Mix. The sample was gel extracted and 4ug were digested with BsaI-HF. The plasmid described above was digested with BsaI-HF for an hour. BsaI was heat inactivated and the digested plasmid was gel extracted before being treated with rSAP for half an hour. Dephoporylated plasmid was cleaned up and 200ng were ligated to the T18 insert at 1:3 ratio with T4 ligase for an hour. Sample was cleaned up and electroporated into freshly prepared electrocompetent pSK34. Approximately 5 million colonies were obtained which were grown up in 200mL LB + Kanamycin + Carbenicillin overnight at 30C. Glycerol stocks were made and stored at -80C. For downstream experiments one tube was fully thawed and subsequently discarded.

4.3 CCNG1 Library cloning

The CCNG1 Library was ordered as part of a 10pM OLS pool and resuspended in 25uL EB. As each interacting pair fit on one oligonucleotide early cloning steps were significantly simplified. The oligonucleotides from the pool were amplified in bulk with oSK229 and oSK232 for all three codon usages. qPCR showed exponential amplification through 16 cycles so a high fidelity PCR was then amplified for 14 cycles in quadruplicate. Samples were pooled and a band of 230bp was run on a 3.5% agarose gel and extracted. Purified sample was diluted 100x and re-amplified for eight cycles in quadruplicate to attach an AscI site (oSK358) and a second BsaI site, 20bp random barcode, and an EcoRI site (oSK359). PCR product was again pooled, run on a 3.5% agarose gel, and extracted for digest with AscI and EcoRI-HF for two hours. Low-copy plasmid pSK33 was purified from 200mL of LB + Kan and 5ug was digested with AscI and EcoRI-HF and rSAP for two hours. The plasmid digest was then run on a 1% agarose gel and the linearized fragment was extracted. 250ng of the linearized plasmid and 1:3 ratio of barcoded OLS product were ligated with T7 ligase for three hours. The ligation product was cleaned up and eluted in 6uL of ddH2O. Electrocompetent NEB 5-alpha cells (NEB C2989K) were transformed with 1uL of the ligation product, cells were recovered for 35 minutes in SOC and plated on large LB + Kan plates. Two million colonies were obtained, and colony PCR showed 23/24 containing the insert. 1.2 million colonies were scraped, diluted to OD 0.02 and grown up in 200mL of LB + Kan for DNA purification.

The T25 insert was cloned as follows: it was amplified from pSK59 with oSK360 and oSK361. The sample was gel purified and digested at 3ug scale sequentially with BspQI and BtsαI for four hours each. 5ug of the plasmid from the previous step was digested with BspQI and Btsαl, for five hours and overnight, respectively, before a 30 minute dephosphorylation with rSAP. The previously purified plasmid and the T25 segment were ligated with T7 ligase for four hours and cleaned up before transformation into freshly prepared electrocompetent pSK34. After electroporation, cells were recovered for 35 minutes and plated onto large LB + Kan + Carb plates. Colonies were scraped, diluted to OD 0.02 in 200mL of LB + Kan + Carb and grown to saturation and plasmid was purified.

The T18-sfGFP insert was cloned as follows: it was amplified from pSK59 with oSK201 and oSK202. The sample was run on a 1% agarose gel and purified and 4ug of it were digested with BsaI-HF for two hours. Likewise, the previously purified plasmid was digested BsaI-HF at 5ug scale for two hours with rSAP, run on a 1% agarose gel and gel extracted. 250ng of plasmid was ligated with the T18-sfGFP insert at a 3:1 ratio with T7 ligase for two hours at room temperature. The sample was cleaned up and electroporated into freshly prepared pSK34. Cells were recovered in SOC for 35 minutes at 37C and plated onto large LB + Kanamycin + Carbenicillin plates. Plates were grown overnight, and all two million cells were scraped diluted to OD 0.02 and grown overnight in LB + Kanamycin + Carbenicillin. Glycerol stocks were made from overnight culture and frozen at -80C. For downstream experiments one tube was fully thawed and subsequently discarded.

4.4 CCmax Library cloning

The CCmax Library was ordered as a 10pM OLS pool and resuspended in 20uL of EB. qPCR with oligonucleotides oSK470 and oSK471 showed exponential exponential amplification through ten cycles, so a high-fidelity PCR was repeated for eight cycles. The amplified product was cleaned up and diluted. Reamplification with qPCR using oSK472 to attach an AscI site and oSK473 to attach a BsaI site, the random DNA barcode and EcoRI site showed exponential amplification through twelve cycles so we performed a high-fidelity PCR for eight cycles with in triplicate. Samples were pooled and run on a 3% agarose gel. A band corresponding to the expected 290bp was extracted, and digested with AscI and EcoRI-HF for two hours at 1ug scale. pSK33 was grown up in 200mL of LB + Kanamycin, purified, and digested at 3ug scale with AscI, EcoRI-HF and rSAP. Digested product was run on a 1% agarose gel and extracted. 250ng of digested pSK33 and a 3:1 ratio of the insert were ligated with T7 ligase for 3 hours. The sample was cleaned up, and 1uL was electroporated into NEB 5-alpha cells (NEB C2989K). Cells were recovered for 35 minutes in SOC media and plated on to LB agar + Kanamycin. Approximately four million transformants were obtained of which 1.2 million were scraped from the plate, and diluted to OD 0.02 in 150mL of LB + Kanamycin and grown up overnight at 37C and DNA was purified.

The T25 insert was cloned as follows: it was amplified from pSK179 with oSK474 and oSK475. The sample was run on a 1.5% agarose gel and then was digested at 4ug scale sequentially with BspQI and Btsαl for four hours each. 5ug of the previously purified plasmid was also digested with BspQI and Btsαl, for three hours and five, respectively, before a 30 minute dephosphorylation with rSAP. The digested plasmid and the T25 insert were ligated with T4 ligase for four hours and cleaned up before transformation into freshly prepared electrocompetent pSK34. After electroporation, cells were recovered for 35 minutes and plated onto LB agar + Kanamycin + Carbenicillin plates. After three successive transformations 500,000 transformants were obtained all of which were scraped from the plates, diluted to OD

0.02, grown up in 200 mL of LB + Kanamycin + Carbenicillin overnight at 37C, and DNA was purified.

The T18 insert was cloned as follows: it was amplified from pSK168 with oSK476 and oSK477. The sample was run on 1% agarose gel, extracted and digested at 3ug scale with BsaI-HF. The previously purified plasmid was digested with BsaI-HF and rSAP for four hours at 5ug scale. Digested product was run on a 1% agarose gel and extracted. 250ng of vector was ligated at 3:1 ratio with the T18 insert with T4 ligase for two hours. The ligation product was purified and electroporated into freshly prepared electrocompetent pSK34. Cells were recovered for 35 minutes in SOC at 37C and plated on LB agar + Kanamycin + Carbenicillin. Plates were scraped diluted to OD 0.02 and grown in 200mL LB + Kanamycin + Carbenicillin to saturation and glycerol stocks were stored at -80C. For downstream experiments one tube was fully thawed and subsequently discarded.

4.5 GFP Library cloning

Each design was synthesized de novo from oligonucleotides ordered from IDT. Ribosome binding sites were synthesized in oSK218-222 and barcodes were attached from oSK217. After cloning into backbone pSK33, we chose 10 colonies from each ribosome binding site, and sequenced their barcodes. Sequenced colonies were inoculated into a deep well plate of LB + Kan + Carb, grown to saturation, pooled and frozen in glycerol stocks at -80C. For downstream experiments one tube was fully thawed and subsequently discarded.

4.6 Individual construct cloning

DNA for the NGB2H system was ordered from Addgene or DNA 2.0. For the bacterial two-hybrid, we used pEB1029 and pEB1030 (Addgene 22066 and 22067)[13]. The pSC101 origin was drawn from pZS-123 (Addgene 26598)[14]. Most of our functional features were drawn from the work of the Voigt lab and iGEM. Inducible promoters for both pTet and pPhlF[3], strong terminators[4], and ribozyme elements upstream of our open reading frames[15] were all synthesized *de novo*. Linkers between *B. Pertussis* proteins and the proteins of interest were drawn from the original B2H with the T18 linker having an GTG extension in the center[1]. sfGFP was drawn from a previously published strain[10]. DNA for all subsequent plasmids was assembled with standard molecular biology techniques, namely Gibson assembly[16] and restriction enzyme digestion/ligation. Individual constructs were sequenced verified before experimental use.

**5. Library mapping:**

All mapping steps used KAPA SYBR Fast qPCR (Kapa Biosciences KK4601), NEBNext Q5 Hotstart HiFi PCR Master Mix (NEB M0543L) for high-fidelity PCR, an Agilent Tapestation with D5000 ScreenTape (Agilent 5067-5583) for DNA quantification, PhiX sequencing control v3 (Illumina FC-110-3001) as a control library and an Illumina Miseq with a v3 600 cycle paired end kit (Illumina TG-142-3003) for sequencing unless otherwise noted.

5.1 CC0 Library mapping

After cloning the CC0 Library proteins and barcodes into pSK33 we sequenced the barcode through the proteins on a MiSeq to provide a mapping function between the two. The amplicon containing the CC0 Library proteins and barcode was amplified with oSK696 and oSK193 or oSK194 for the different replicate barcodings. This attached P5, P7 and a Nextera lowplex index to allow demultiplexing of the barcoding replicates. qPCR showed exponential amplification for 15 cycles for both samples, so a high-fidelity PCR was repeated for 12 cycles in triplicate. Samples were pooled and a band corresponding to the expected size of 391bp was extracted from a 2% agarose gel. The sample concentrations were measured on an Agilent Tapestation 2200 and found to be monodispersed with a length of 405bp. The samples were mixed in equimolar amounts with 15% PhiX control, diluted to 14pM and loaded into the Illumina Miseq kit. We used three separate custom primers, oSK696 for the forward read, oSK323 for the reverse read and oSK324 for the index read. We obtained 26,359,427 reads which mapped to 137,213 unique barcodes with a correct X or Y protein for the first barcoding replicate and 96,129 unique barcodes with a correct X or Y protein in the second barcoding replicate.

5.2 CC1 Library mapping

The CC1 Library mapping step was performed similar to the CC0 Library mapping. After cloning the CC1 Library and random barcode in pSK33, we amplified constructs with either oSK691, oSK692 or oSK693 and oSK193 to attach Illumina adaptor P5 and a Nextera lowplex I7 and Illumina adaptor P7 respectively. qPCR showed exponential amplification through 14 cycles. PCR was repeated with high-fidelity polymerase, KAPA HiFi HotStart ReadyMix (KAPA KK2602) and the samples were cleaned up and pooled. Concentration of the pooled sample was measured with KAPA Library Quantification Kit Illumina Systems (Kapa Biosystems KK4824) and found to be 26nM. This was diluted to 12pM, mixed with 5% phiX control, and run on a Miseq with a pool of oSK709, oSK710 and oSK711 for the forward read, oSK324 for the index read and oSK323 for the reverse read. 8,060,843 reads passed filter which mapped 1,166,860 unique barcodes mapping to one or both proteins from the CC1 Library.

5.3 CCNG1 Library mapping

Similar to the CC0 Library mapping, after cloning the barcoded CCNG1 Library into pSK33 we sequenced the barcode through the proteins with a MiSeq. The amplicon containing the CCNG1 Library was amplified using oligonucleotides oSK366 and oSK193 to attach P5, Nextera lowplex index N702 and P7. qPCR showed exponential amplification through eleven cycles so samples were amplified for high-fidelity PCR for nine cycles in triplicate. Samples were pooled and a band corresponding to the expected size of 376bp was gel extracted from a 1.5% agarose gel. The sample concentration was quantified on an Agilent Tapestation 2200 and found to be monodispersed and of approximately 376bp. The sample was diluted to 20pM and mixed with 10% PhiX control, loaded into the Miseq kit, and sequenced with custom primers oSK367 for the forward read, oSK323 for the reverse read and oSK324 for the index read. 27,255,659 reads passed filter which mapped 1,121,668 unique barcodes to one or both proteins from the CCNG1 Library.

5.4 CCmax Library mapping

Similar to the CC0 Library mapping, after cloning the CCmax Library into pSK33, sequenced through the proteins and the barcode with a Miseq. The amplicon containing the CCmax proteins and barcode was amplified with oSK513 and oSK193. This was first done with qPCR which showed exponential amplification for 1ng through 15 cycles. High-fidelity PCR was repeated in triplicate for nine cycles, samples were pooled and run on a 1.5% agarose gel and a band corresponding to the expected 380bp was extracted. Sample concentration was quantified with an Agilent Tapestation 2200 and was found to be monodispersed and approximately 384bp. The sample was mixed with 10% PhiX control and 18pM was loaded into an Miseq kit. To the MiSeq Kit we added custom primers oSK514 for the forward read, oSK323 for the reverse read and oSK324 for the index read. 20,710,707, reads passed filter which mapped to 1,629,936 unique barcodes mapping to one or both proteins in the CCmax Library.

5.5 Mapping script

To map DNA barcodes to a unique interaction we used a custom Makefile that chained several programs from the BBTools suite[17] with our own custom script. From the raw paired fastq files we used BBduk (v38.32) to trim adapter sequences and remove contaminants. Forward and reverse reads were then merged with BBmerge (v38.32) with strictness set to maxloose. Merged reads with levenshtein distance three or less were then condensed with Starcode (v1.3) to remove sequencing errors. From the condensed reads, barcodes were mapped with a custom python script. Briefly, this script called the last twenty bases per sequence the barcode and discarded those barcodes within levenshtein distance one of each other. It then removed those barcodes that had proteins that were more than 5% different from each other, under the assumption that these were contaminants. The variants were then mapped to the DNA corresponding to the protein coding regions using BBMap (v38.32) of the X and Y proteins sequentially. Sequencing data that mapped with no errors to a reference sequence of X or Y were then joined together by barcode and this text file was used to identify barcodes from barcode sequencing steps.

#### 6. NGB2H experimental conditions and validation

6.1 CC0 Library NGB2H assay

Glycerol stocks of both barcodings of the CC0 Library were thawed, and 100uL were grown up overnight in 100mL EZ Rich Defined Media (Teknova M2105) with Kanamycin (Teknova K2125) and Carbenicillin (Teknova C2130). After overnight growth 1mL of the GFP Library was added to the 100mL of CC0 Library culture and 1mL of the GFP Library was added to a fresh culture of 100mL EZ Rich Defined Medium + Carbenicillin + Kanamycin with 25ng/mL anhydrotetracycline (aTC), 1uM 2,4-Diacylphlorolglucinol (DAPG) and 100uM Isopropyl B-D-1-thiogalactopyranoside (IPTG) in biological replicates for each barcoding. Flasks were incubated in a 37C shaker for six hours. Samples were pulled at 6h and placed on an ice slurry for 15 minutes, spun down for nucleic acid extraction and flash frozen. As reported in the main text we obtained high quality verification of the CC0 Library from its internal controls. Briefly, we obtained 57,541 barcodes providing quantitative measurements of interaction strength for all 256 protein pairs. The assay was highly replicable with biological replicates having similar Interaction Scores (Pearson's $r$ > 0.98, $p < 10^{-15}$) (Figure 1C). Different codon usages showed consistent Interaction scores with all usages correlating with Pearson's $r$ > 0.92 and $p < 10^{-15}$ (Figure S5). The CC0 Library has a strong correlation between the primary and reciprocal orientations (Pearson's $r$ = 0.92, $p < 10^{-15}$) (Figure 1E). The Interaction Scores for constructs with indels was much less than full length perfect constructs (Figure S6). When compared to the published Tms of the CC0 Library the

Interaction Scores correlated with Pearson's *r* > 0.73, $p < 10^{-15}$ (Figure S7). Finally, when the assay is replicated with an independent re-barcoding and re-cloning of the library, we found very strong replication with the previous CC0 Library's Interaction Scores (Pearson's *r* > 0.98, $p < 10^{-15}$) (Figure 1F).

6.2 CC1 Library NGB2H assay

Glycerol stocks were thawed and we grew the CC1 Library overnight in EZ Rich Defined Medium + Kanamycin + Carbenicillin. We mixed 1:100 of the GFP Library with the CC1 Library and diluted it to OD 0.001 in a fresh EZ Rich media + antibiotics with 5ng aTC and 5uM DAPG at 30C to induce the library. The library was grown in a time-course experiment, where we took samples of RNA and DNA at 0h, 0.5h, 1h, 2h and 4h which were spun down and flash frozen for nucleic acid preparation. In total we linked 385,078 different barcodes 400 different PPIs. Our internal controls validated that the library performed as expected. After normalization to a constitutive GFP Library, Interaction Scores were minimal at the beginning of the assay, but they increased monotonically over four hours of induction (Figure S8A). In addition, comparison of six different codon usages in the CC1 Library showed high replicability (*r* > 0.89, $p < 10^{-15}$, Figure S8B). Finally, compared to full length constructs, indels had a markedly reduced Interaction Score (Figure S8C) and the reciprocal orientation of each protein were similar (*r* > 0.85, $p < 10^{-15}$, Figure S8D).

6.3 CCNG1 Library NGB2H assay

After overnight growth in EZ Rich media + Carbenicillin + Kanamycin , we mixed in a constitutive GFP Library to the CCNG1 Library at a 1:100 ratio and took RNA and DNA at 0h. We induced a 1:100 dilution of the library for 6h with 25ng/mL aTC, 15uM DAPG and 100uM IPTG in EZ Rich media + antibiotics at 37C in biological replicates. We obtained 76 million reads across 164,778 barcodes which gave quality data on 8,073 interactions. Our internal controls for the CCNG1 Library behaved as expected. The Interaction Score of constructs in the CCNG1 Library strongly correlated between biological replicates (Pearson's *r* > 0.95, $p < 10^{-15}$, Figure S10A); the CC0 Library subset of the CCNG1 Library correlated strongly with published Tms (Pearson's *r* > 0.79, $p < 10^{-15}$, Figure S10B) and with our previous experiments of the CC0 Library (Pearson's *r* > 0.94 , $p < 10^{-15}$, Figure S16). Furthermore, different codon usages for the same construct gave similar Interaction Scores (Pearson's *r* > 0.75, $p < 10^{-15}$, Figure S10C), and indels had significantly Interaction scores than full length constructs (Figure S10D).

6.4 CCmax Library NGB2H assay

After overnight growth in EZ Rich media + Kanamycin + Carbenicillin, we mixed the CCmax Library with the GFP Library in 100:1 ratio. We diluted the library by a 1:100 ratio in fresh EZ Rich media with 15ng/mL aTC, 5uM DAPG and 100uM IPTG in biological replicates for 6h at 37C. We obtained 346,733 barcodes mapping to 17,731 constructs and collected high quality data on 17,320 interactions. We found that biological replicates strongly correlated (Pearson's *r* = 0.973, $p < 10^{-15}$, Figure S15A), as did different codon usages with (Pearson's *r* > 0.92, $p < 10^{-15}$, Figure S15B). As expected full length perfect constructs had higher Interaction Scores than those containing indels in the X or Y protein (Figure S15D). We again included the CC0 Library found that the published Tms correlated with our Interaction Score with (Pearson's *r* = 0.876, $p < 10^{-15}$, Figure S15C), and that correlated well with the CC0 proteins in other libraries (Pearson's

$r > 0.84$, $p < 10^{-15}$, Figure S16). Finally, the reciprocal orientation of the proteins in the CCmax Library largely agrees with their primary one, with (Pearson's $r = 0.835$, $p < 10^{-15}$, Figure S15E).

**7. Barcode sequencing preparation**

All nucleic acid preparation for barcode sequencing was done with Qiagen kits. For RNA prep we used RNeasy Midi (Qiagen 74106) or RNeasy Minipreps (Qiagen 75144) with on column DNase digestion (Qiagen 79254) and concentrated with RNeasy MinElute Cleanup kit (Qiagen 74204). DNA was prepped with QIAprep spin Minipreps (Qiagen 27106) or Plasmid Plus Maxiprep (Qiagen 12963), though DNA cleanup and gel extraction was performed with Zymo kits (Zymo D4014 and Zymo D4008). As for previous steps, we used NEBNext Q5 Hotstart HiFi PCR Master Mix (NEB M0543L) for high-fidelity PCR and KAPA SYBR Fast 2x Master Mix (Kapa Biotechnology KK4601) for qPCR. All samples were quantified with Agilent D5000 Screentape (Agilent 5067-5582).

<u>7.1 RNA Barcode Sequencing Preparation</u>

Cell pellets containing either 1 x 10^10 cells or 1 x 10^9 cells were thawed and purified with midi or mini scale respectively with on column DNase digestion. RNA was concentrated and purified RNA was subject to primer specific reverse transcription (oSK193-oSK198 and oSK210-oSK215) to create cDNA of transcripts containing DNA barcodes. We used the reverse transcription step to attach Nextera lowplex I7 indexes and P7 Illumina sequence adaptors to our barcodes which allowed multiplexed sequencing of different times and conditions. Reverse transcription was performed with Superscript IV (ThermoFisher 18090050) with the following changes to the manufacturer's protocol. Instead of 5ug of RNA we used 22.5ug of RNA, concentrated to 11uL, the reverse transcription step at 55C was allowed to go for an hour rather than fifteen minutes, and 1uL RNAase A (Qiagen 19101) was spiked in with RNase H for twenty minutes. After reverse transcription samples were amplified with qPCR with oligonucleotides oSK199 and oSK200 to attach Illumina sequencing adapter P5. We compared the cDNAs with no-RT controls to check for DNA contamination in the RNA which invariably showed less than 1:1000 ratio of DNA to RNA. The qPCR showed exponential amplification through 20 cycles. We then repeated the PCR in triplicate for 12-15 cycles. Replicate PCRs were pooled, run on a 3% agarose gel and extracted. DNA concentration was measured on an Agilent Tapestation 2200 with D1000 screentape (Agilent 5067-5582), and equimolar fractions pooled with DNA barcodes.

<u>7.2 DNA Barcode Sequencing Preparation</u>

Cell pellets containing either 10mL or 50mL of culture were thawed and plasmid DNA was extracted with mini or maxi scale respectively. Barcodes were amplified with qPCR using oSK199 and oSK193-198 or oSK210-215 to attach Illumina sequencing adaptor P5, and a Nextera lowplex I7 index and Illumina sequencing adaptor P7. We used qPCR to guide exponential amplification of the barcodes which normally showed exponential amplification through 13-16 cycles. We repeated the PCRs for 10-12 cycles in triplicate. Replicate PCR samples were pooled, run on a 3% agarose gel, and extracted. DNA concentration was measured on an Agilent Tapestation 2200 and equimolar fractions were pooled with the RNA barcodes.

#### 8. NGB2H small scale results:

8.1 Plate reader measurements:

Strains used in plate reader assays were grown up overnight in MOPS EZ Rich Defined Media (Teknova M2105) with kanamycin (Teknova K2125) and carbenicillin (Teknova C2130) in a 37C degree shaker. The next evening these cultures were diluted 1:100 in 100uL fresh MOPS EZ Rich Defined Media with kanamycin, carbenicillin and 100uM IPTG and the indicated inducers in a 96 well, flat bottom plate (Corning 0720090). The plate was then incubated in a Tecan M1000 Plate Reader at 37C overnight. Optical Density (OD600) and GFP fluorescence (excitation 488nm, emission 508nm) were taken every half an hour after 3 minutes of 1mm orbital shaking. Data was collected for a minimum of fourteen hours but normally reached saturation by eight hours.

8.2 Single construct optimization and benchmarking

We found the NGB2H system behaved as expected with sfGFP fluorescence dependent on a pair of interacting proteins and both anhydrotetracycline (aTC) to induce pTet and 1,4-Diacylphloroglucinol (DAPG) to induce pPhlF (Figure S1A). Lacking either inducer or assaying a pair of non-interacting proteins yielded only low levels of sfGFP fluorescence. We reasoned that basal sfGFP expression would correspond to more noise in our multiplexed experiments, as non-interacting barcodes would constitute a larger proportion of all sequenced barcodes. Thus, we empirically optimized the signal to noise ratio of induction by testing a panel of Jun/Fos constructs where pPhlF and pTet had varying ribosome binding sites and sfGFP was driven by several pLac variants. We selected a construct that gave 96x signal of induced/uninduced sfGFP fluorescence, called pSK59 (Figure S1B). Although our overall signal strength was weaker with the PhlF RBS variant we selected compared to some constructs assayed, there was extremely little sfGFP fluorescence in our uninduced sample (Figure S1C) which we reasoned would lead to higher signal in the multiplexed assay.

To evaluate the quantitative range of our assay we analyzed a previously published set of coiled-coils with $K_d$s ranging from $10^{-6}$ to $10^{-10}$ M,[18] that as well as an additional construct with an inferred $K_d < 10^{-6}$. Measuring sfGFP fluorescence, we found that our system can detect weak interactions, as low as $10^{-6}$ (Figure S1D), however, it lacks the power to discriminate between medium ($10^{-7}$) to high ($10^{-10}$) $K_d$s. In contrast with a previous study using the standard *B. pertussis* adenylate cyclase two-hybrid system, our modified system enables us to achieve quantitative measurements in agreement with published $K_d$s[19].

8.3 CC1 Library results

The designs from the CC1 Library were expected to be orthogonal within each backbone subset. The expected orthogonal design of the coiled-coils was largely recapitulated in our results (Figure 9A), with the P#s having only one strong interaction, P3/P12, which was unexpected. The P#mS and P#SC# backbones also exhibited the expected orthogonal pattern within their subsets. As P3mS-P8mS and P5SC1-P6SC2 contain the same interfacial residues as their P# counterparts we expected them to cross react with the corresponding P# partners. We found this to be the case with the P#mS having the same reaction pattern (Figure 9A bottom, grey) with the P# set as with itself. The P#SC# set also cross reacted as expected but had the unexpected reaction of P6SC#/P7 (Figure 9A, bottom orange).

We were curious if our interaction profiles matched what would be expected from our designs. We defined a favorable electrostatic interaction as an E- or G-position having a Glu or Lys forming a salt bridge with a Lys or Glu, respectively, in the corresponding position on the partner protein. Likewise, we defined an Ile/Ile or Asn/Asn pair as having isoleucines or asparagines in the A-position of a given heptad for both proteins. We found that our strong interactions highly favored having all eight possible salt bridges, as nearly all interactions with eight salt-bridges have a higher Interaction score than all other interactions (Figure 9B, top). We found the identity of the residue at position A to be less determinative. Though most constructs with a high Interaction score had four pairs of Ile/Ile or Asn/Asn at position A, there were many constructs with four pairs of Ile/Ile or Asn/Asn that did not have a high Interaction score (Figure 9B, lower). Taken as a whole, these designs largely functioned as expected: Glu/Lys pairings were far more preferable to Glu/Glu or Lys/Lys pairings and Ile/Ile and Asn/Asn pairs, rather than Ile/Asn pairs, were necessary but not sufficient to create an interaction.

8.4 Effects of backbone variation in the CCNG1 Library

Though the B-, C- and F-position residues are thought to modulate binding affinity, the CCNG1 Library is the first large dataset that systematically tests the effects of different backbones. A subset of the CCNG1 Library contains the interfacial residues of the CC0 Library on six different backbones, which vary from nearly exclusively small non-polar residues to nearly exclusively large charged residues (Figure S13A). To understand how the different backbones affect specificity we divided interactions into on-target and off-target groups where the on-target group had an interface with published Tms greater than 60C (Figure S13B). On-target interactions invariably had high interaction scores and no significant difference was noted between the backbones, though the bH backbone did have higher variance than the rest. Off-target interactions, however, unexpectedly showed that the original backbone had lower Interaction scores than the other backbones. To investigate this further we compared each protein, as determined by the interfacial residues, against the previously published Tm (Figure S13C). We found that all backbones except the original were strongly shifted to higher interaction scores. Although further tests are needed to understand why there is a global shift to higher interaction scores with other backbones--especially given that highly helical amino acids such as alanine are expected to produce the strongest coiled-coil interactions--one possible hypothesis is that the presence of glutamine in these backbones can facilitate low strength off-target interactions.

When broken down into categories of, small and large underestimates and overestimates, a clear pattern emerged that every backbone had an over representation of interaction scores that were higher than the original backbone, and many were much higher than expected (>1 interaction score more). Conversely, the bA backbone had only a handful of interactions below the expected strength and none that were strongly so. Taken together this suggests the need for care when using backbones with polar residues as unpredicted effects may occur, particularly at the expected lower range.

**Supplementary References**

1. Karimova, G., Pidoux, J., Ullmann, a & Ladant, D. A bacterial two-hybrid system based on a reconstituted signal transduction pathway. *Proc. Natl. Acad. Sci. U. S. A.* **95**, 5752–5756 (1998).

2. Battesti, A. & Bouveret, E. The bacterial two-hybrid system based on adenylate cyclase reconstitution in Escherichia coli. *Methods* **58**, 325–334 (2012).
3. Stanton, B. C. *et al.* Genomic mining of prokaryotic repressors for orthogonal logic gates. *Nat. Chem. Biol.* **10**, 99–105 (2014).
4. Chen, Y.-J. *et al.* Characterization of 582 natural and synthetic terminators and quantification of their design constraints. *Nat. Methods* **10**, 659–64 (2013).
5. Badran, A. H. *et al.* Continuous evolution of Bacillus thuringiensis toxins overcomes insect resistance. *Nature* **533**, 58–63 (2016).
6. Kuhlman, T., Zhang, Z., Saier, M. H. & Hwa, T. Combinatorial transcriptional control of the lactose operon of Escherichia coli. *Proc. Natl. Acad. Sci. U. S. A.* **104**, 6043–8 (2007).
7. Crooks, R. O., Lathbridge, A., Panek, A. S. & Mason, J. M. Computational Prediction and Design for Creating Iteratively Larger Heterospecific Coiled Coil Sets. *Biochemistry* **56**, 1573–1584 (2017).
8. Gradišar, H. & Jerala, R. De novo design of orthogonal peptide pairs forming parallel coiled-coil heterodimers. *J. Pept. Sci.* **17**, 100–106 (2011).
9. Islam, S. *et al.* Characterization of the single-cell transcriptional landscape by highly multiplex RNA-seq. *Genome Res.* **21**, 1160–1167 (2011).
10. Kosuri, S. *et al.* Composability of regulatory sequences controlling transcription and translation in \textlessi\textgreaterEscherichia coli\textless/i\textgreater. *Proc. Natl. Acad. Sci.* **110**, 14024–14029 (2013).
11. Potapov, V., Kaplan, J. B. & Keating, A. E. Data-Driven Prediction and Design of bZIP Coiled-Coil Interactions. *PLoS Comput. Biol.* **11**, 1–28 (2015).
12. Brodnik, A., Jovičić, V., Palangetić, M. & Silađi, D. Construction of orthogonal CC-sets. *Informatica* **43**, (2019).
13. Battesti, A. & Bouveret, E. Improvement of bacterial two-hybrid vectors for detection of fusion proteins and transfer to pBAD-tandem affinity purification, calmodulin binding peptide, or 6-histidine tag vectors. *Proteomics* **8**, 4768–4771 (2008).
14. Cox, R., Dunlop, M. J. & Elowitz, M. B. A synthetic three-color scaffold for monitoring genetic regulation and noise. *J. Biol. Eng.* **4**, 10 (2010).
15. Lou, C., Stanton, B., Chen, Y. J., Munsky, B. & Voigt, C. A. Ribozyme-based insulator parts buffer synthetic circuits from genetic context. *Nat. Biotechnol.* **30**, 1137–1142 (2012).
16. Gibson, D. G. *et al.* Enzymatic assembly of DNA molecules up to several hundred kilobases. *Nat. Methods* **6**, 343–345 (2009).
17. Brian Bushnell. BBMap.
18. Fletcher, J. M. *et al.* A basis set of de novo coiled-Coil peptide oligomers for rational protein design and synthetic biology. *ACS Synth. Biol.* **1**, 240–250 (2012).
19. Smith, A. J., Thomas, F., Shoemark, D., Woolfson, D. N. & Savery, N. J. Guiding Biomolecular Interactions in Cells Using *de Novo* Protein–Protein Interfaces. *ACS Synth. Biol.* **8**, 1284–1293 (2019).

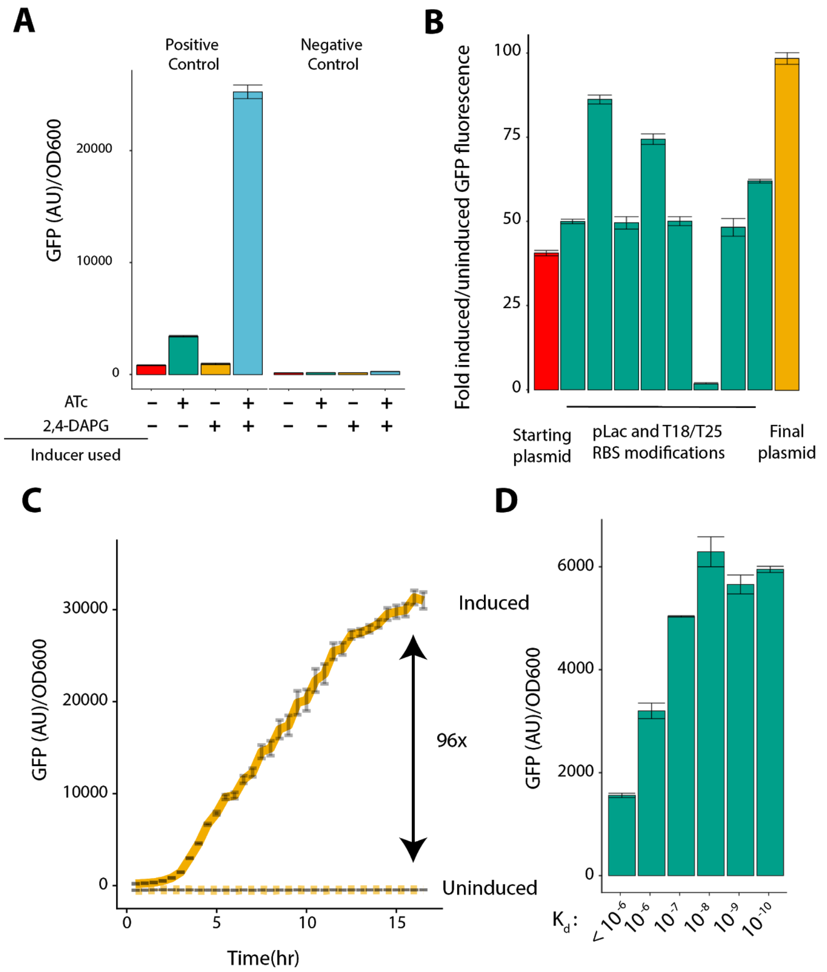


**Figure S1) Optimization and tuning of the NGB2H system:** A) High GFP expression requires positively interacting proteins and both inducers, ATc and DAPG. Error bars are standard deviation of three technical replicates B-C) Optimizations of the promoters and RBSes to find the maximal signal to noise ratio between induced and uninduced samples. B) The ratio of (Induced GFP/OD fluorescence)/(Uninduced GFP/OD fluorescence) for Jun/Fos constructs with RBS and promoter variations. Error bars represent standard deviation of three replicates. C) The Final plasmid induced or uninduced over 16hr. Samples were taken every 30 minutes. Error bars represent standard deviation of three replicates D) A panel of previously characterized proteins shows GFP/OD depends on $K_d$ with maximal expression occurring at $K_d$'s stronger than $10^{-7}$ M. Error bars represent standard deviation of three replicates.

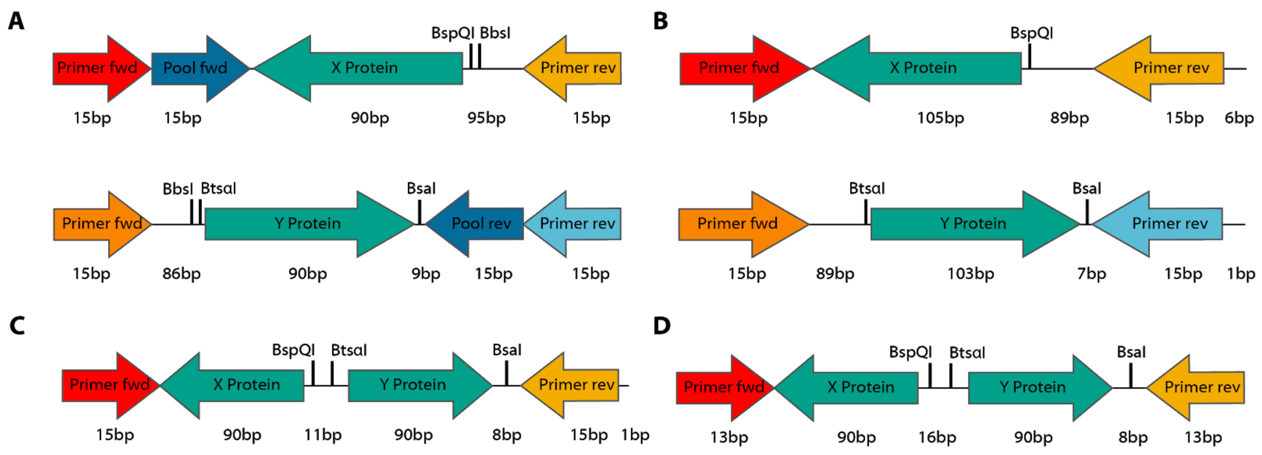


**Figure S2) Design of OLS oligonucleotides for libraries used in this work:** With minor variation each oligonucleotide consists of primers to amplify it out of the OLS pool, the coding sequence and several restriction enzyme sites. Numbers below the constructs represent how CC0 Library oligonucleotides are divided into those with the X protein and those with the Y protein. B) CC1 Library oligonucleotides are divided into those coding the X protein and those coding the Y protein. C) CCNG1 Library oligonucleotides contain both the X and Y protein on a single oligonucleotide. D) CCmax Library oligonucleotides contain both the X and Y protein on a single oligonucleotide.

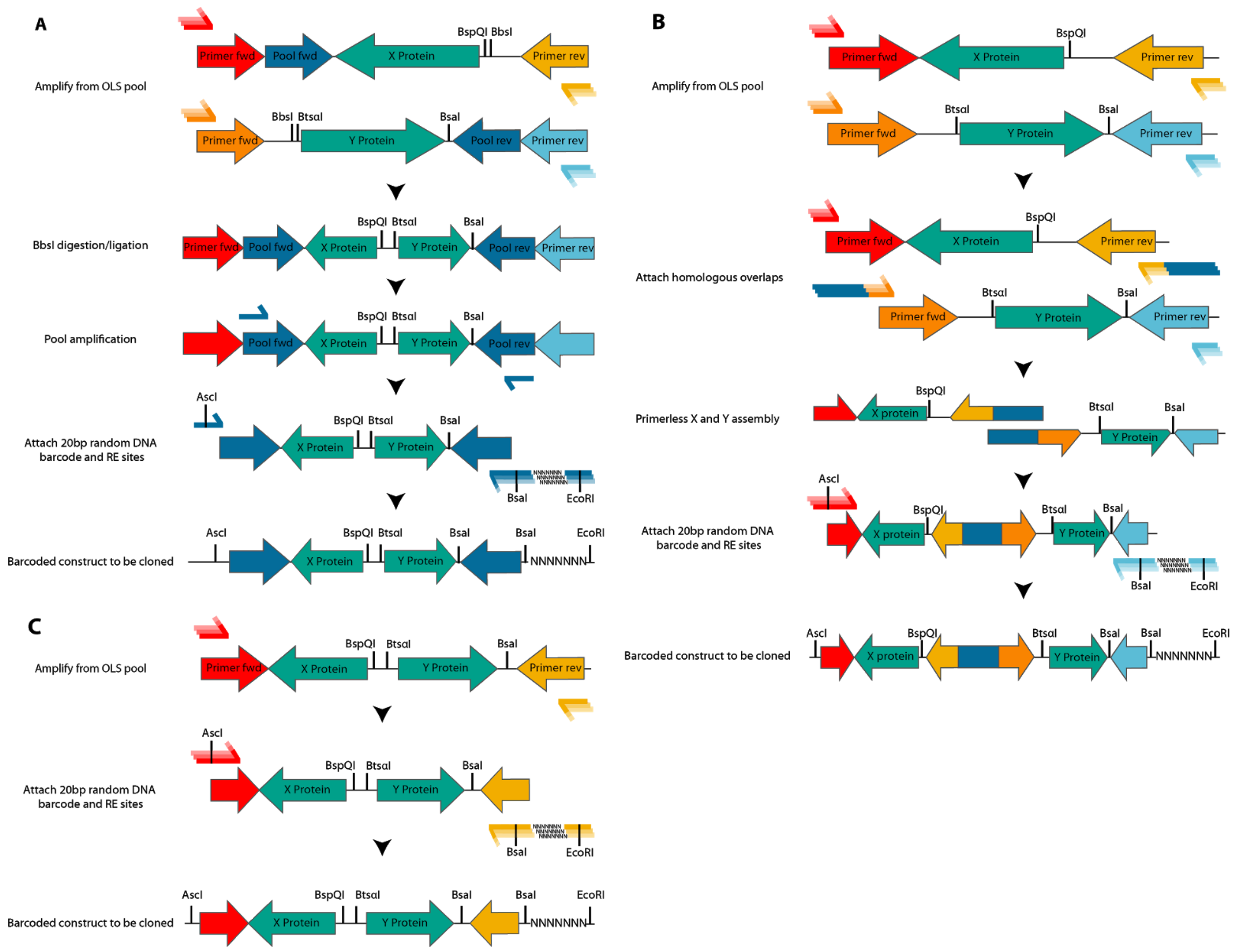


**Figure S3) Cloning from OLS oligonucleotides to barcoded X and Y constructs:** A) CC0 Library barcoding schema. Five OLS pools for both the X and Y oligonucleotides are amplified with OLS primers. All oligonucleotides are digested with BbsI and ligated in pairs, before amplification with pool primers and mixing. Finally, restriction enzyme sites for cloning into the vector and the barcode are attached. B) CC1 Library barcoding schema. After amplification of the OLS pool, matching overlaps are attached which are stitched together with overlap PCR. Finally, the barcode and restriction enzyme sites for cloning into the vector are attached. C) CCNG1 and CCmax Library barcoding schema. Oligonucleotides are amplified from the pool and then restriction enzyme sites for cloning into the vector and barcode are attached.

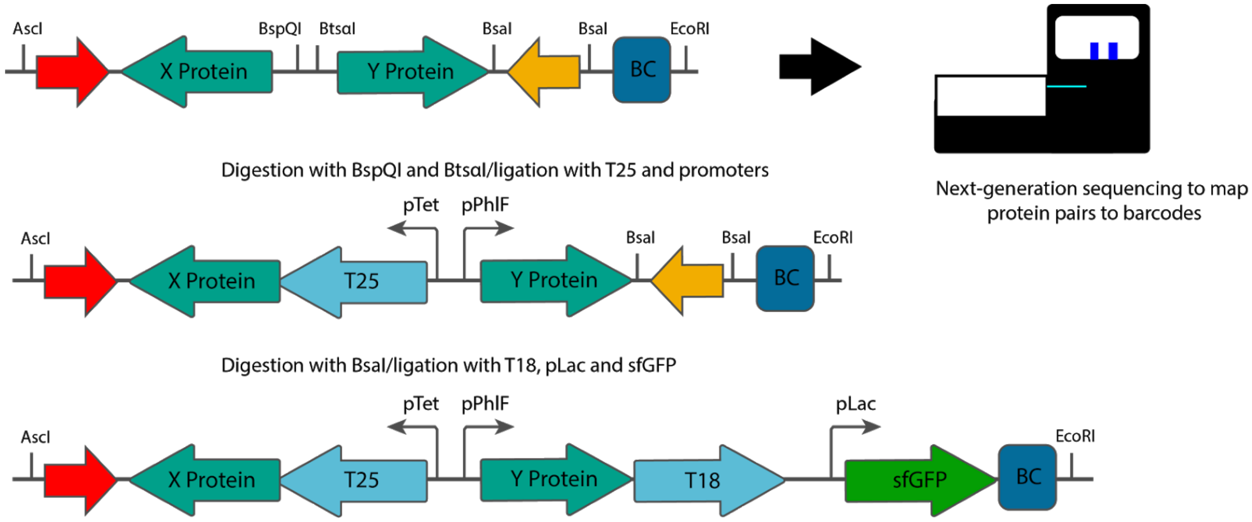


**Figure S4) Cloning scheme of the NGB2H system after barcoding:** After barcoding, the X and Y proteins are sequenced through the barcode using an Illumina MiSeq to identify each barcode's corresponding protein pair. After mapping, the T25 section and inducible promoters are cloned into the mapped plasmid. After the T25 section is cloned, the T18 section with sfGFP is cloned into the T25 containing plasmid.

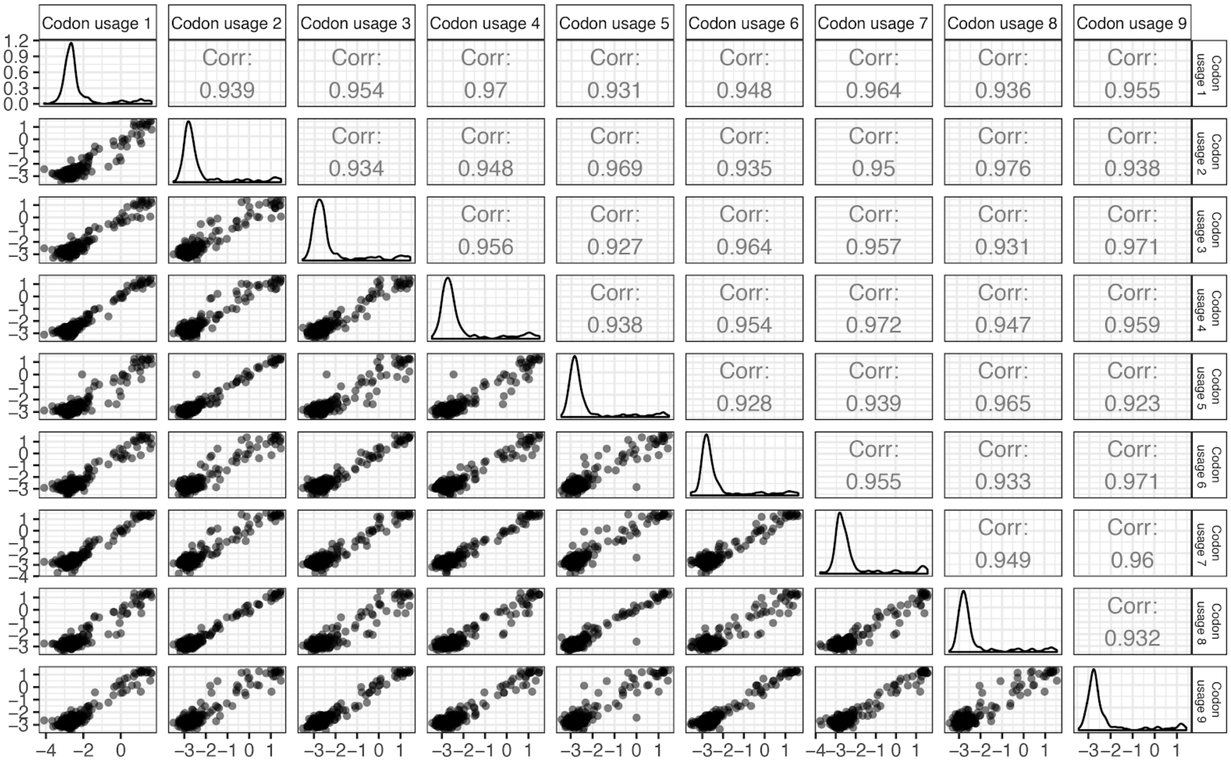


**Figure S5) Different codon usages of the CC0 Library:** Different codon usages in the CC0 Library produce similar Interaction Scores. All nine codon usages from the CC0 Library show high replicability with Pearson's *r* > 0.92 in all pairwise interactions and a mean replicability of *r* > 0.949.

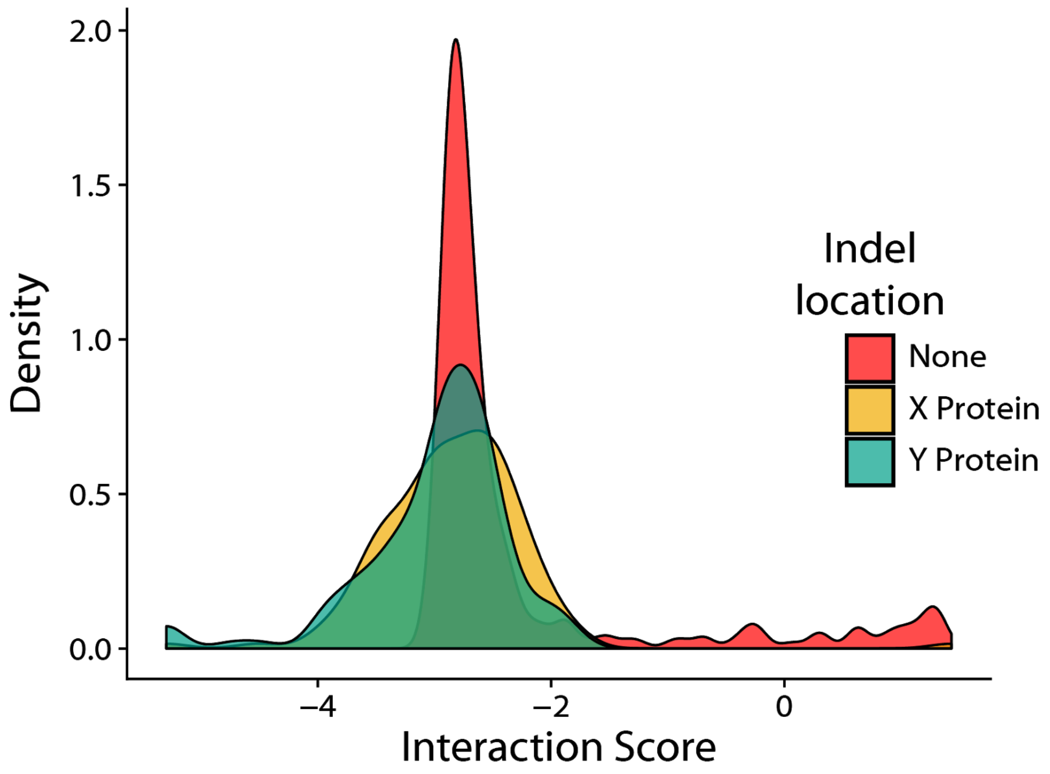


**Figure S6) Indels in the CC0 Library have lower interaction scores than correct sequences:** Constructs with insertions or deletions in the X or Y protein invariably have an Interaction Score of less than -1.8. Constructs without indels, however have some Interaction scores as great as .8.

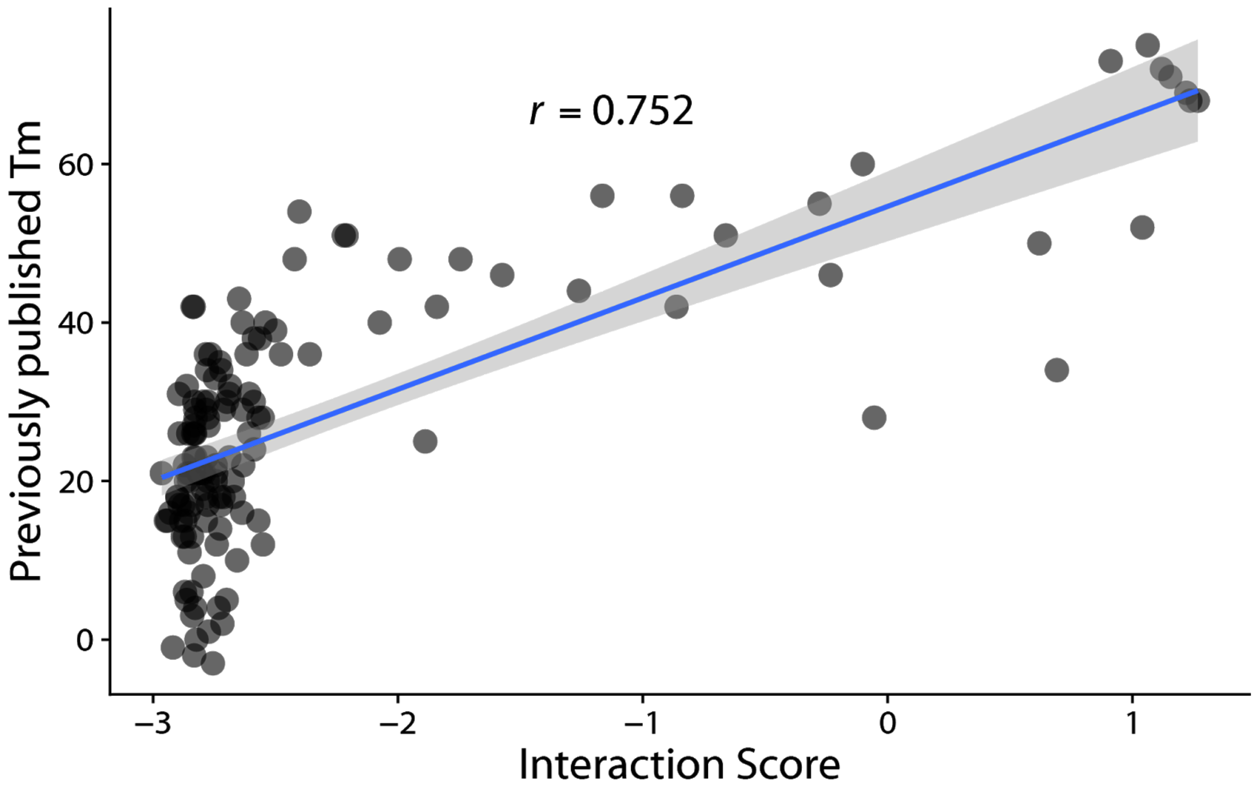


**Figure S7) CC0 Library Interaction Scores versus previously published Tms:** The Tms measured by circular dichroism correlate well with the Interaction Score measured in the CC0 Library with Pearson's *r* > 0.75.  Tms > 40C were well distinguished by the NGB2H assay. Blue line represents a linear model fit to the data, with standard error as the gray shading.

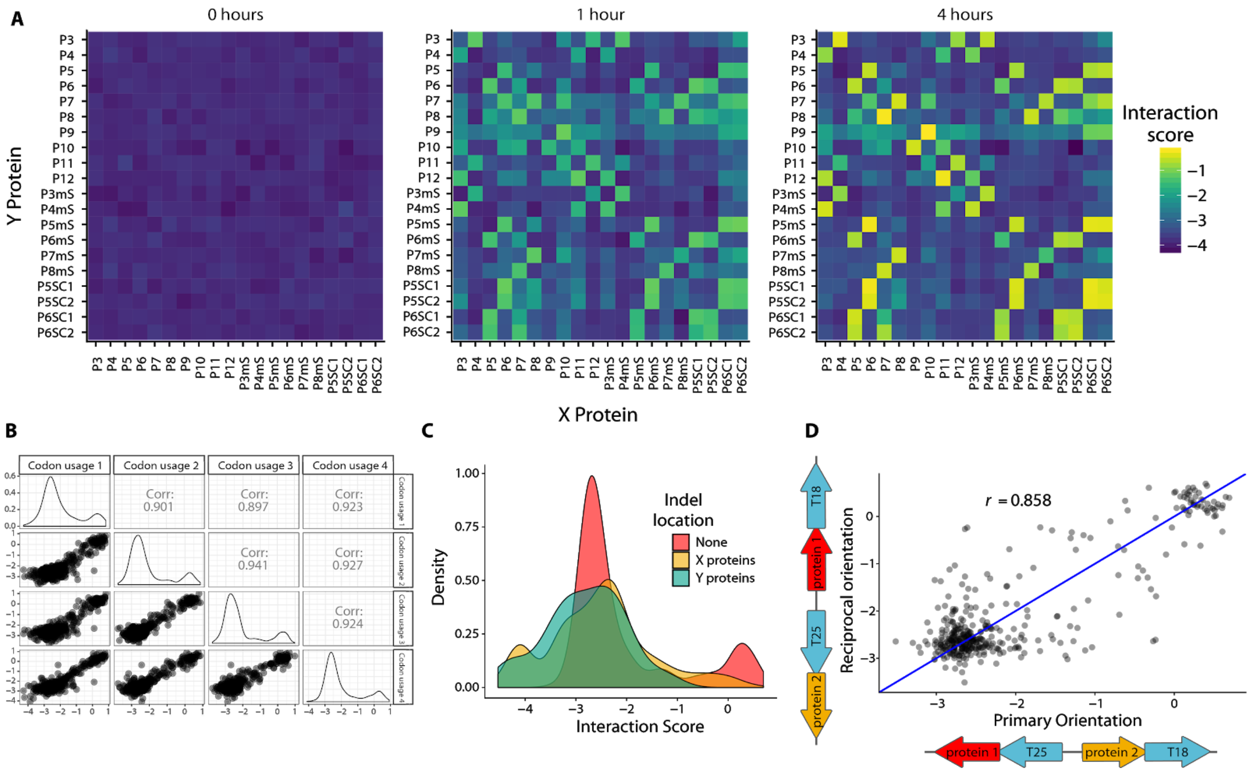


**Figure S8) CC1 Library internal controls:** A) A timecourse experiment of the CC1 Library shows increasing Interaction Scores over four hours, with the strongest signal coming at four hours. Interaction scores were normalized across time with the constitutive GFP Library. B) Different codon usages for the CC1 Library replicate with Pearson's R > 0.89 for all pairwise interactions and a mean of Pearson's *r* = 0.918 C) The Interaction Score for constructs with indels in the CC1 Library is lower than for those without indels. D) The CC1 Library constructs give similar (*r* > 0.85) Interaction Scores for protein pairs attached

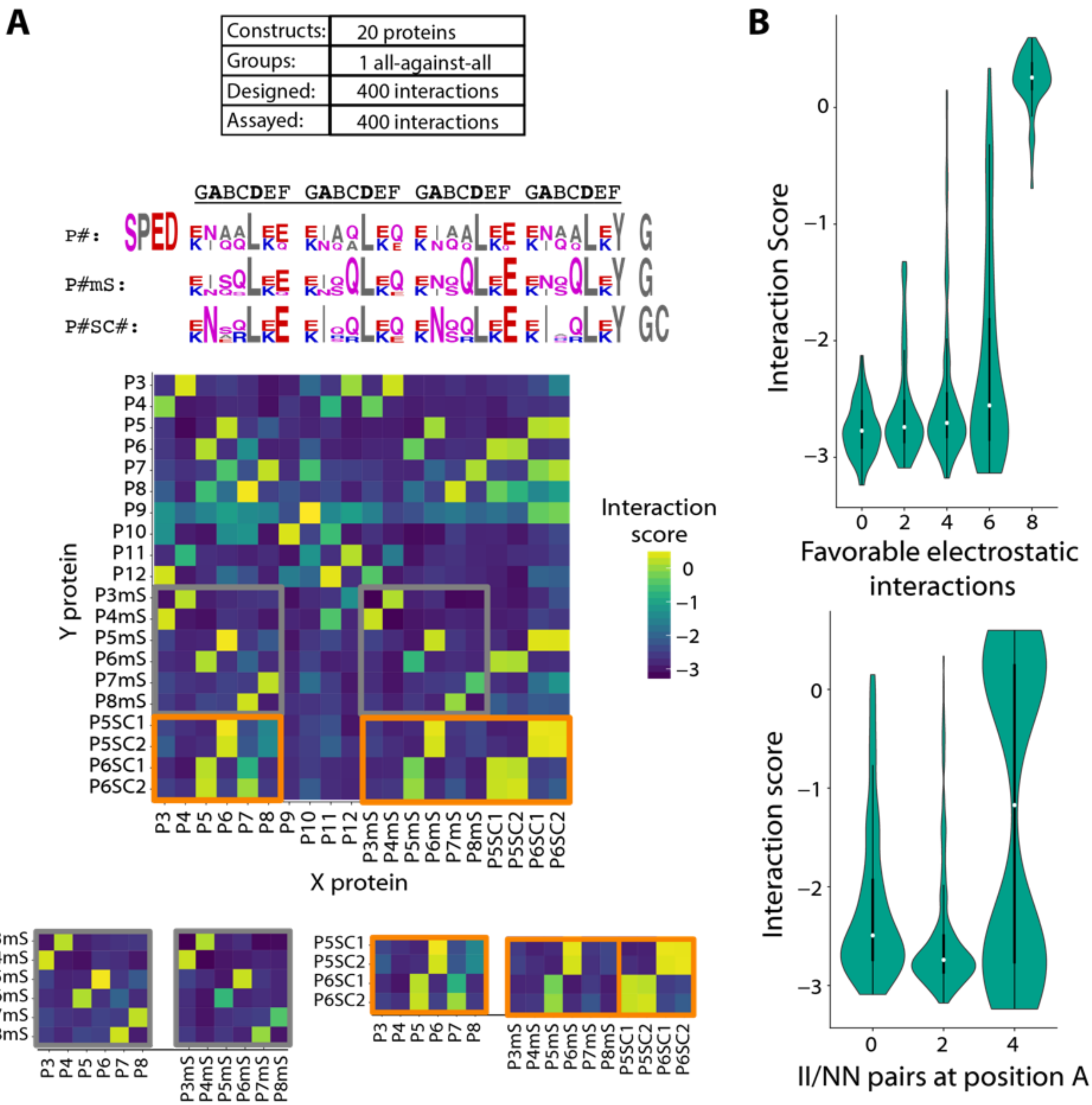


**Figure S9) Isolation of orthogonal coiled-coils and their determinants from the CC1 Library.** A) (Upper) Expected outcome for the CC1 Library and logo illustrating the library diversity. Residues are colored by their functional group Red: positively charged, Blue: negatively charged, purple: polar, grey: non-polar or other (Lower) Interaction scores for entire CC1 Library. The inset in gray shows similarities between library P# and P#mS library members. The inset in orange shows similarities between P#, P#mS and P#SC#. B) (Upper) The Interaction score is highly dependent on the number of electrostatic interactions. White dot represents the median. (Lower) Interaction score is dependent on the residue pairing in the A position, but this is not determinative.

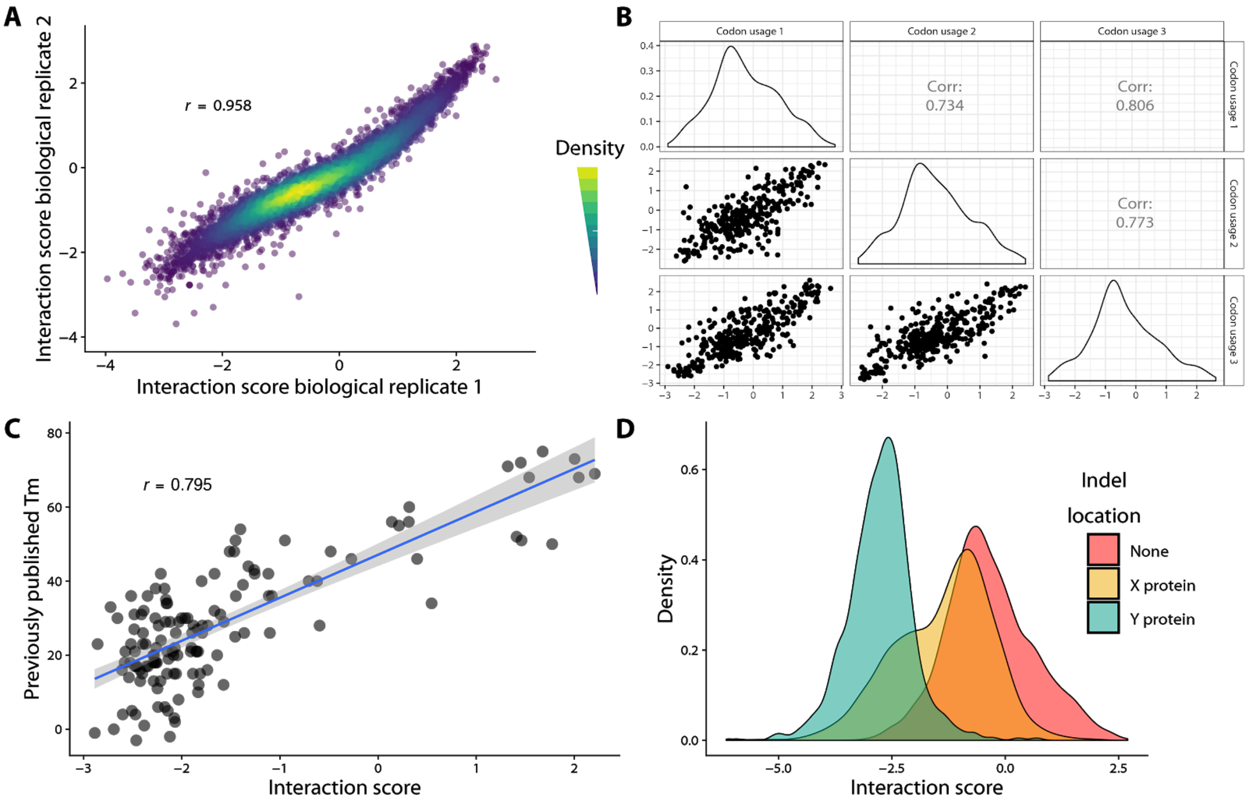


**Figure S10) CCNG1 Library internal controls:** A) The CCNG1 Library replicates in biological replicates with Pearson's $r > 0.95$ ($p < 10^{-15}$). B) Different codon usages in the CCNG1 Library correlate with each other with $r > 0.73$ ($p < 10^{-15}$) for all pairwise comparisons (mean = 0.77). For this analysis only constructs with ten barcodes were used. C) The CC0 Library was included in the CCNG1 Library, and correlates with previously published Tms with Pearson's $r > 0.79$ ($p < 10^{-15}$). Blue line represents a linear model fit to the Interaction scores; grey shading represents the standard errorD) Indels in the CCNG1 Library show decreased Interaction scores compared to constructs without indels.

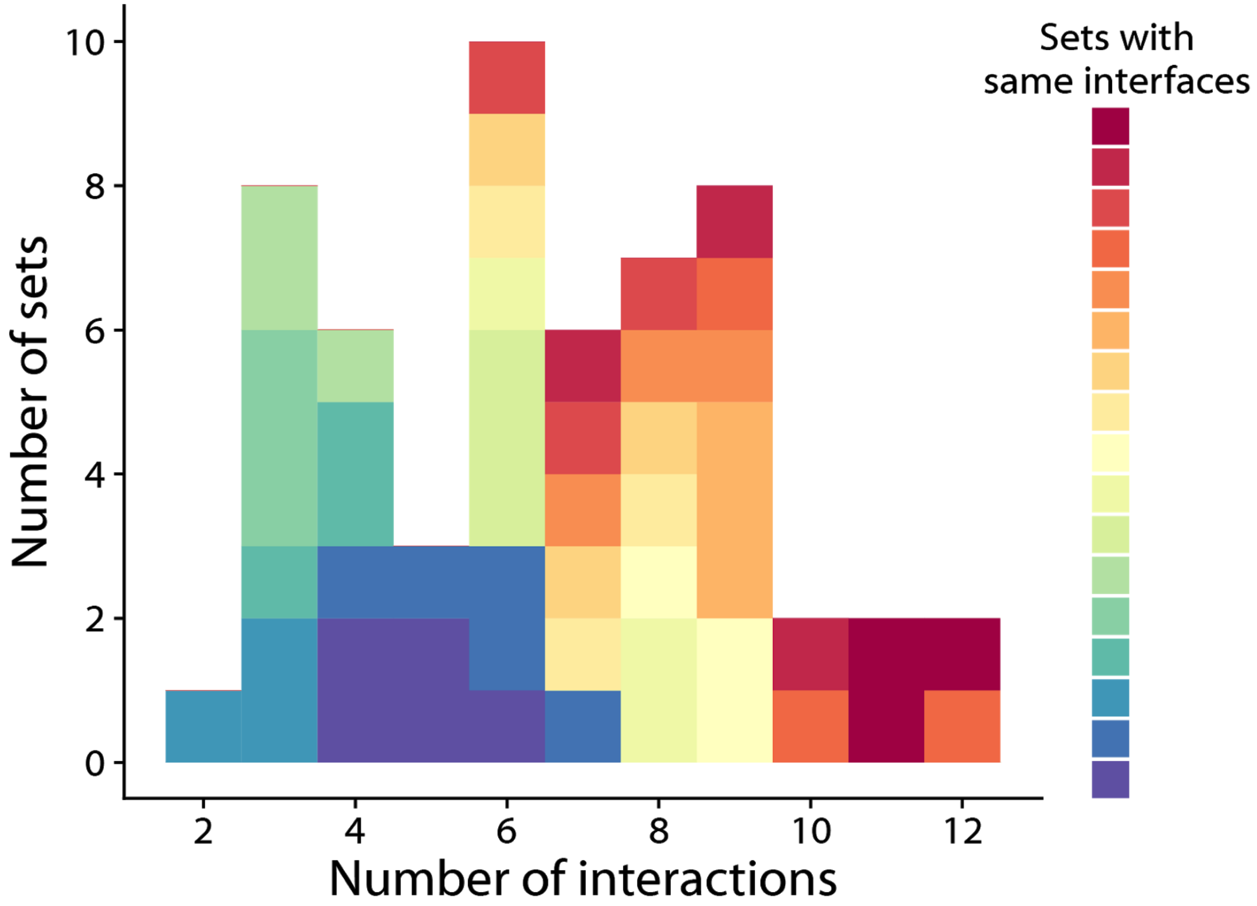


**Figure S11) Number of orthogonal interactions by sets with different backbones in the CCNG1 Library:** Sets with different backbones but the same interfacial residues generally contained similar numbers of orthogonal interactions.

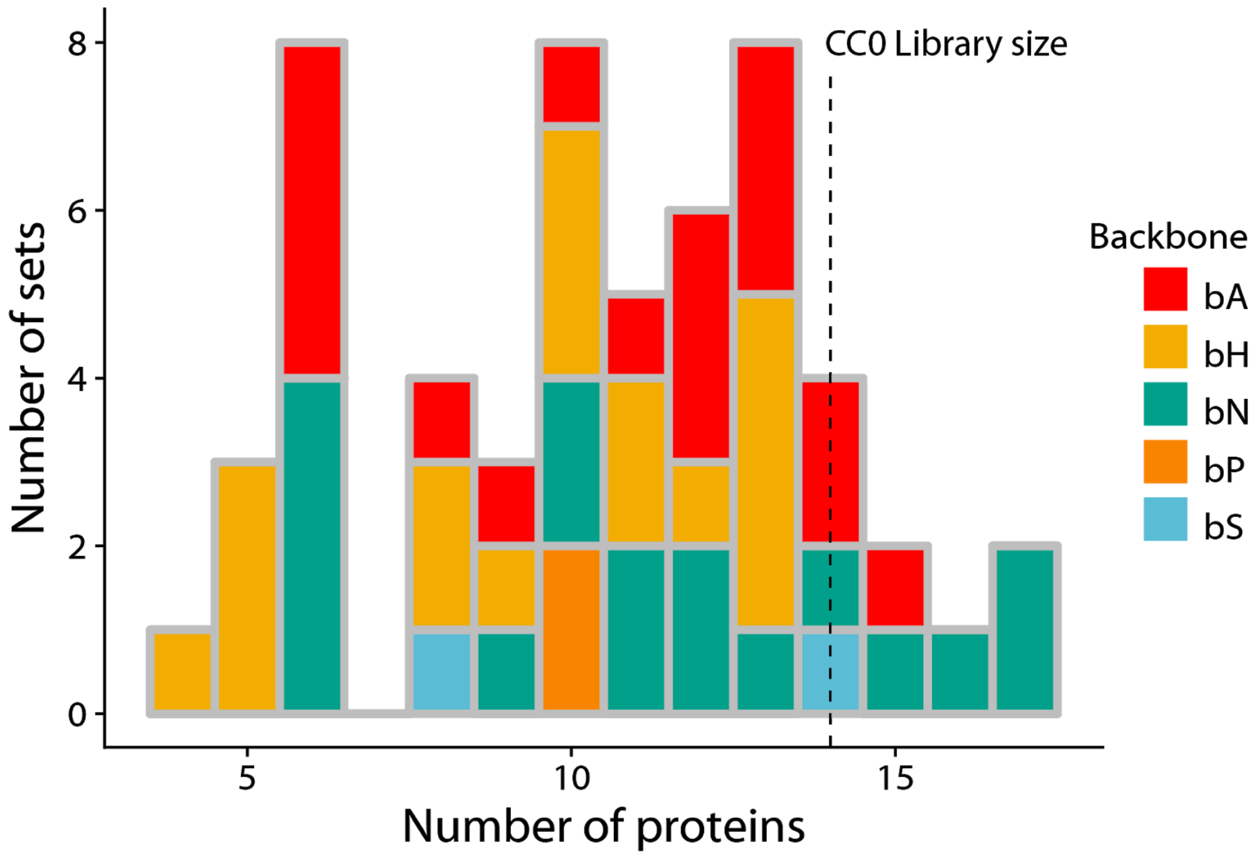


**Figure S12) CCNG1 Library number of proteins per orthogonal set:** Sets in the CCNG1 Library contained between four and seventeen distinct proteins. Five of these contain more than the fourteen in the CC0 Library. Different backbones contained the same interfacial patterns.

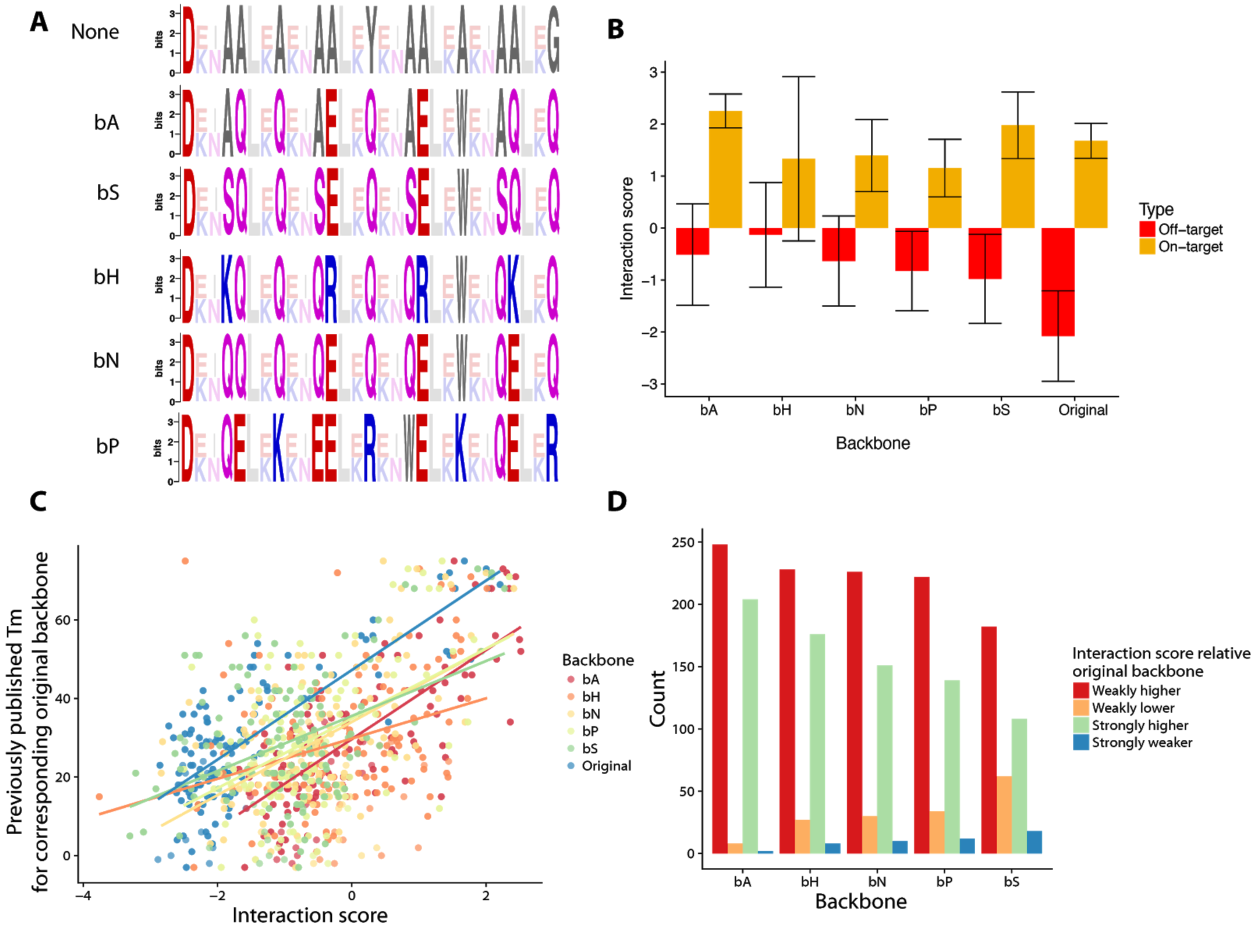


**Figure S13) Effect of variation at the b, c, and f-positions:** A) Sequence logos representing the different backbones. Amino acids are colored according to their type. Red: negative, Blue: positive, Purple: polar, Grey: non-polar. Non-transparent residues are backbone residues. B) Mean Interaction scores for different backbones. On-target is defined as the eight interactions with predicted Tm = 72C with bCipa; off-target interactions are all other interactions. Error bars are standard deviation. C) The Interaction scores from each backbone compared to the Tm of proteins that share the same interfacial residues D) Counts of Interactions scores for different backbones above or below the Interaction score for the original backbone. Strongly higher/lower is defined as an interaction score greater than ±1.

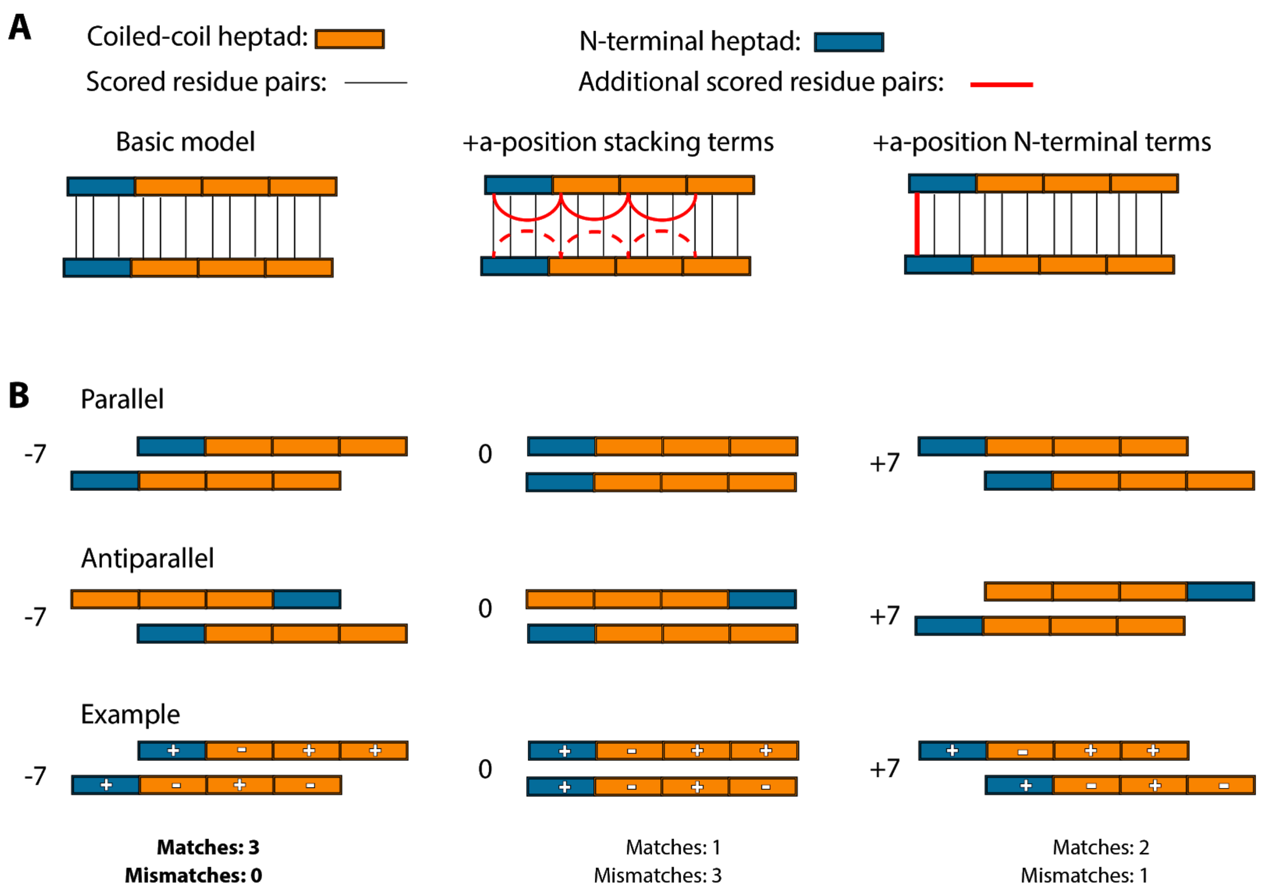


**Figure S14) Schematic of heptad shifting:** A) Different model variations tried for iCipa. The basic model only scores a-, e- and g-positions. The +a position stacking terms model scores consecutive residues in the a-position while the +a-position N-terminal terms model includes separate weights for the first a-position. B) All iCipa candidates score interactions with heptad shifting, that is moving up or down seven residues in an interaction. From left to right shows progressive heptad shifts of the bottom coiled-coil with respect to the top coiled coil for both parallel and antiparallel coiled-coils. (Bottom row) As an example illustrating how heptad shifting is scored, each heptad is given a plus sign or a minus sign, the combination of which is considered a match. In the -7 position all three heptads match giving a high score. In the 0 and 7 positions the heptads have fewer matches and more mismatches so the -7 position would be chosen as the orientation to score. Note though, iCipa calculates individual residues rather than entire heptads at a time.

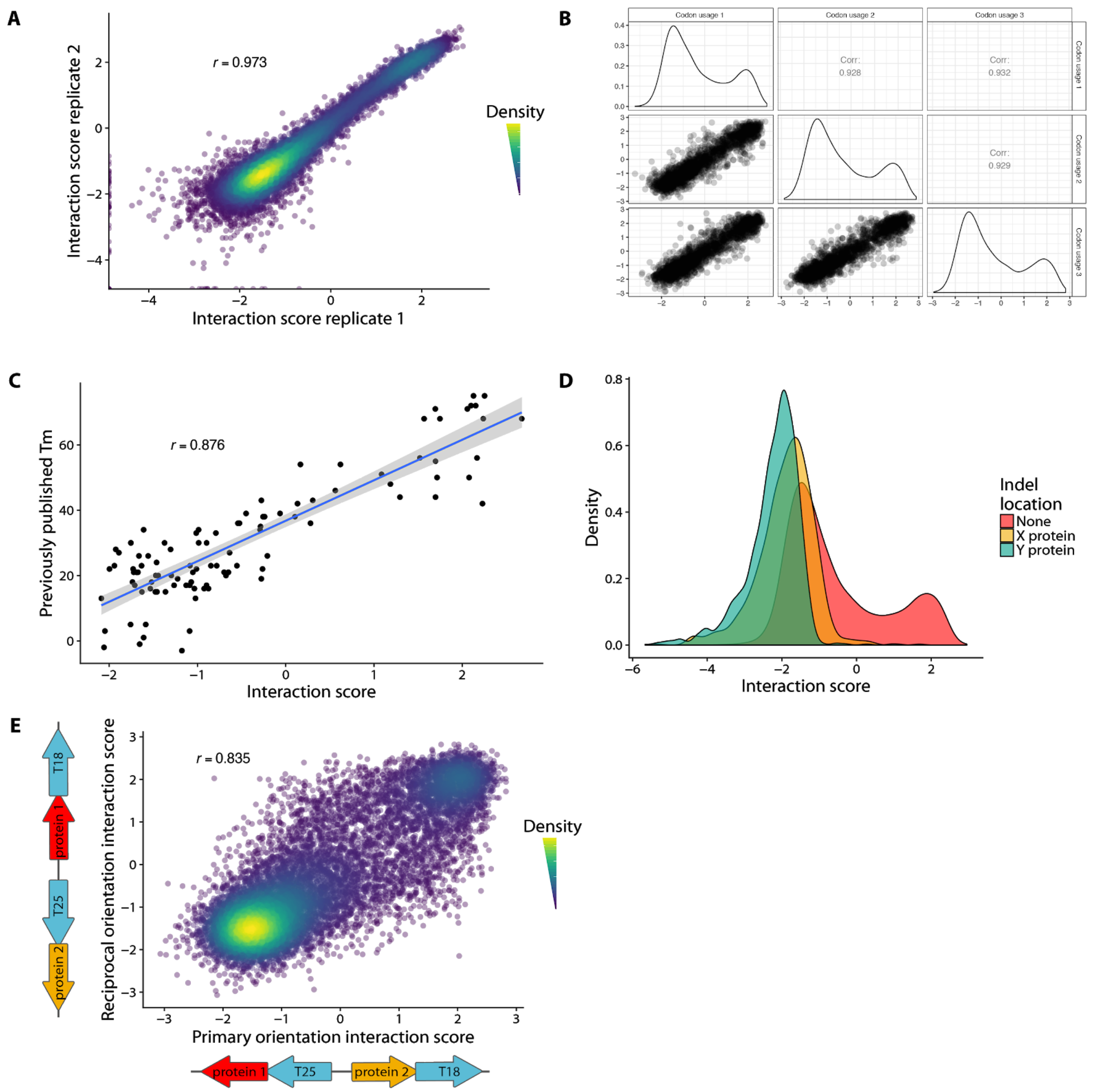


**Figure S15) CCmax Library internal controls:** A) Interaction scores of variants in the CCmax Library correlate strongly between biological replicates (Pearson's *r* > 0.97, $p < 10^{-15}$ ). B) Different codon usages of the CCmax Library have similar Interaction scores with Pearson's *r* > 0.92 and $p < 10^{-15}$ for all pairwise comparisons. C) The CCmax Library contained the CC0 Library. When our Interaction scores are compared to the previously published Tms they correlate well with Pearson's *r* > 0.87, $p < 10^{-15}$ . D) Correct constructs from the CCmax Library have a higher interaction score than those produced with indels. E) The reciprocal orientations of the CCmax Library have similar Interaction scores, and correlate with Pearson's *r* > 0.83, $p < 10^{-15}$ .

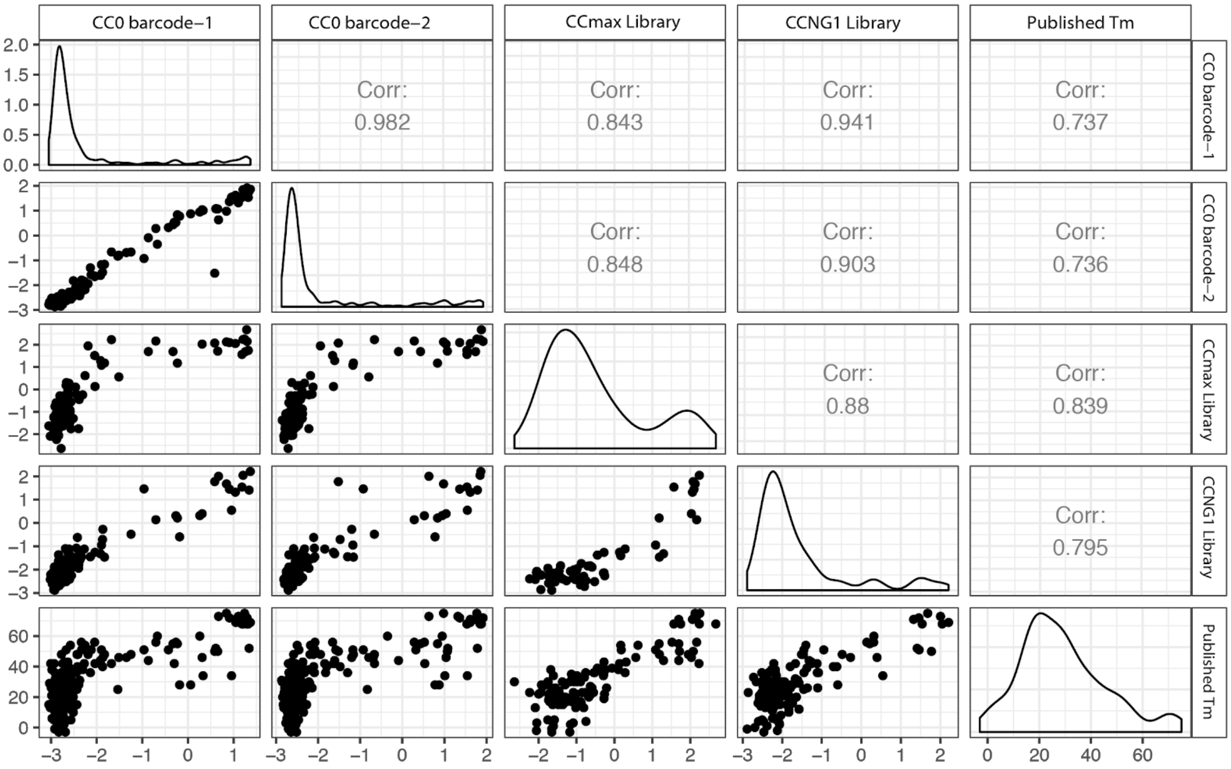


**Figure S16) Correlation of the CC0 Library proteins between different libraries:** The CC0 Library was a subset of all libraries except CC1. Comparing how it performed in all libraries shows strong agreement between sets with Pearson's $r > 0.84$, $p < 10^{-15}$ between all libraries and Pearson's $r > 0.73$, $p < 10^{-15}$ for all libraries with the previously published Tms.

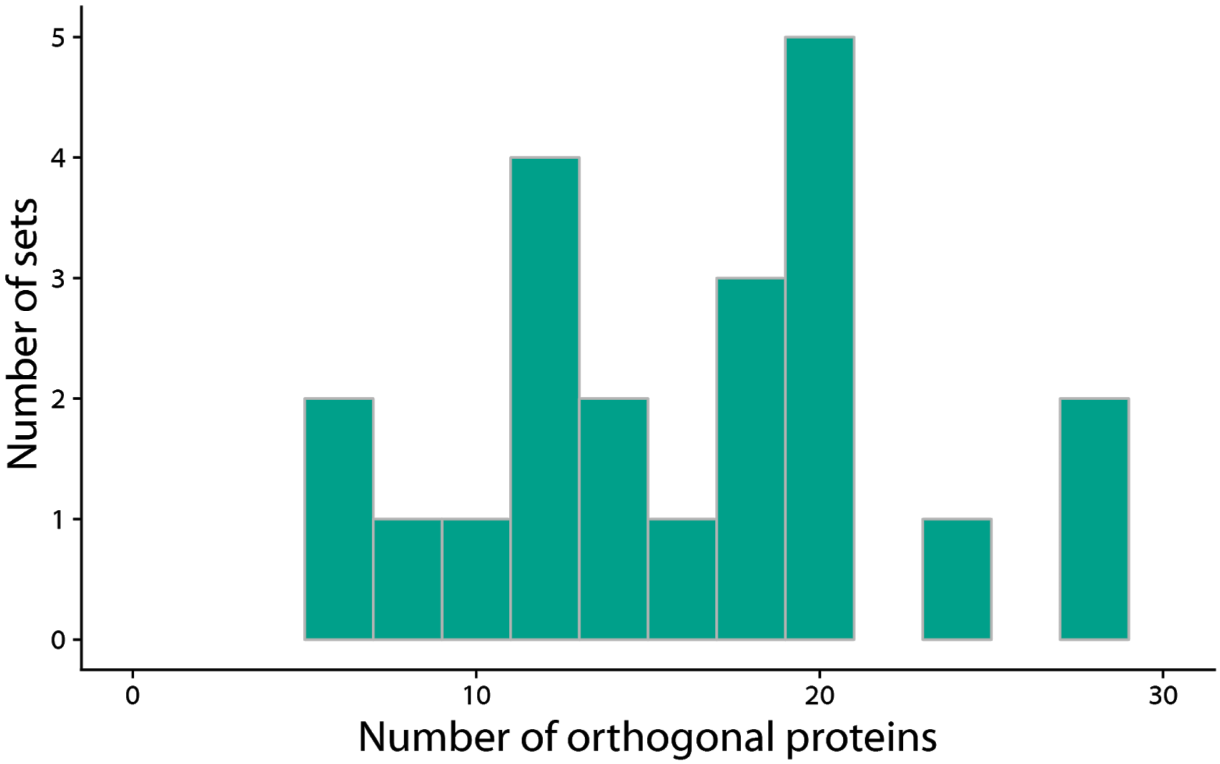


**Figure S17) Number of orthogonal proteins per set:** The CCmax Library had orthogonal sets that contained the most orthogonal proteins of any group of orthogonal proteins to date. Sets contained between 6 and 28 proteins or between 36 and 784 total interactions.

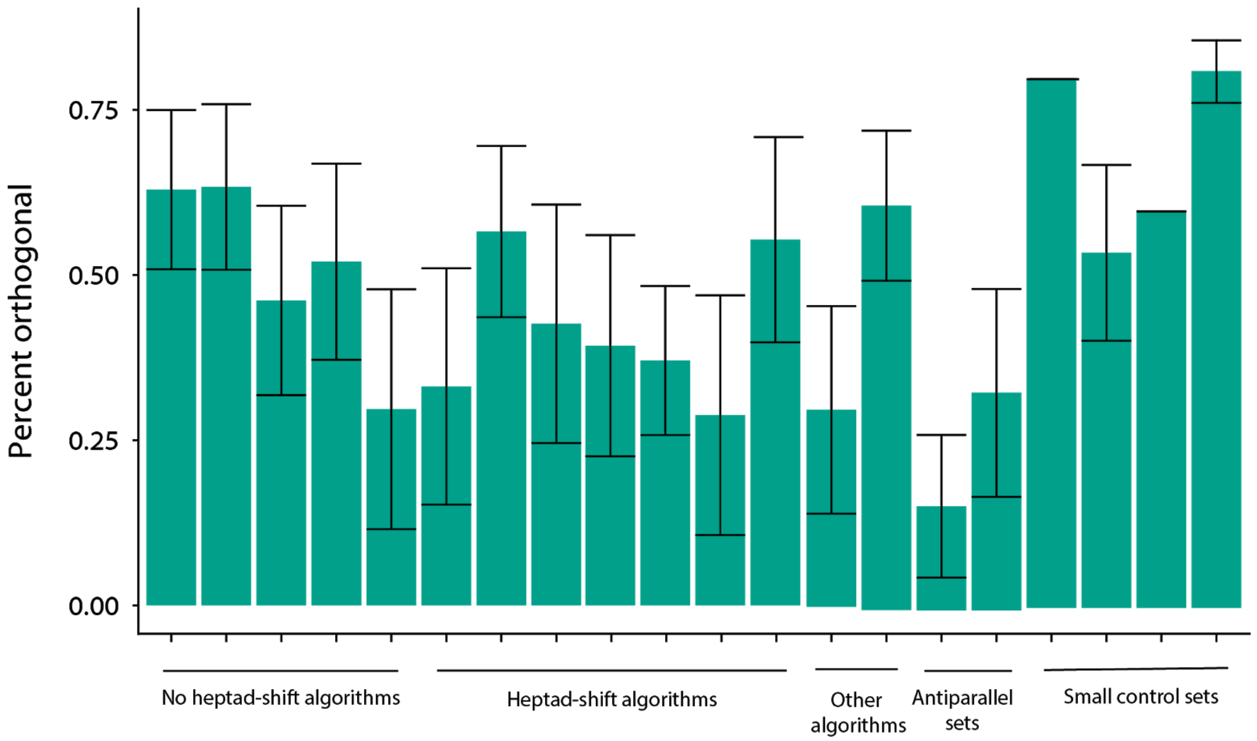


**Figure S18) Ability to predict orthogonality compared across algorithms:** Sets of proteins from the CCmax Library that were designed with different algorithms were randomly subsampled to subsets of ten proteins and the largest orthogonal group was identified. Subsampling was repeated 500 times. Error bars are standard deviation.

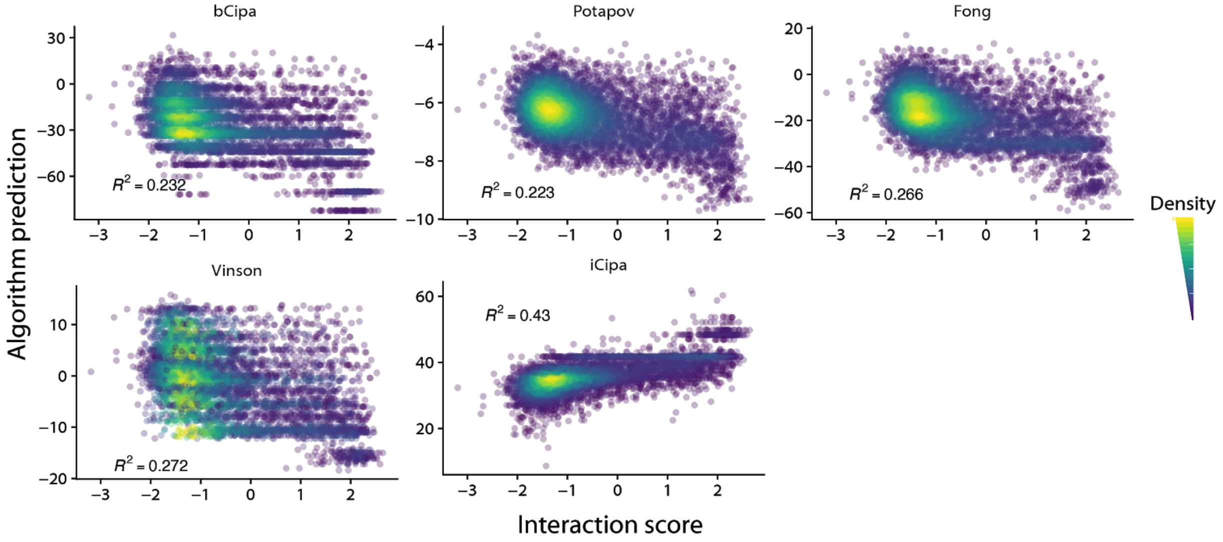


**Figure S19) CCmax Library's agreement with previous models:** Interaction scores from the CCmax Library correlate poorly with previous models. All previous models predict Interaction scores with a coefficient of determination less than 0.28, but iCipa predicts Interaction scores with $R^2 = 0.43$.
